# Jamestown Canyon virus rapidly adapts to mosquito cells through multiple M segment mutations

**DOI:** 10.64898/2026.06.22.733793

**Authors:** Sarah Dysinger, Tamanna Srivastava, Sara Cherry, Paul Bates

## Abstract

Jamestown Canyon virus (JCV) is a mosquito-borne orthobunyavirus with an unusually broad host and vector range. Despite increasing mosquito-to-human spillover, the viral determinants governing host adaptation remain poorly defined. We examined changes in JCV replication during serial passage in mosquito cells and sought to link adaptive changes in viral fitness to specific genetic mutations. In mosquito-derived C6/36 cells, JCV exhibited a distinct “lag-burst” phenotype in which viral replication remained nearly undetectable for 10 days before abruptly increasing. Strikingly, following reinfection of fresh C6/36 cells, JCV that had been passaged once in mosquito cells exhibited immediate, robust replication with no detectable lag phase. Sequencing before and after passage identified multiple M segment mutations associated with enhanced replication. Using a plasmid-based reverse genetics system, individual mutations were introduced into recombinant JCV and evaluated for their effects on replication in mosquito cells. All tested mutations independently enhanced replication efficiency, demonstrating that adaptation can arise through multiple independent genetic pathways. However, no individual mutation fully reproduced the phenotype acquired naturally through mosquito cell passage. Together, these findings demonstrate that JCV rapidly adapts to mosquito cells under minimal selective pressure and highlight the potential for emergence of increasingly well-adapted viral variants.

## 1. INTRODUCTION

Jamestown Canyon virus (JCV), a mosquito-borne orthobunyavirus in the family *Peribunyaviridae*, was originally isolated from mosquitoes in 1961 in Jamestown, Colorado (1, 2). Subsequently, JCV was found to be widely distributed across the northern United States and Canada (3, 4). As an arbovirus, JCV circulates between mosquito vectors and vertebrate hosts, with white-tailed deer serving as the primary amplification host (3, 5, 6). However, evidence of infection has also been documented in numerous incidental hosts, including caribou, dogs, cattle, horses, foxes, and polar bears, among others (7–9). Serological studies indicate that exposure is common in both reservoir and incidental hosts, with regional seroprevalence in deer exceeding 80%. Up to 50% of humans in endemic areas show evidence of prior exposure but humans are considered dead-end hosts (10). Unlike other arboviruses that are restricted to one or two competent mosquito vectors, JCV has been detected in more than 33 mosquito species spanning multiple genera, and several species are thought to sustain year-round transmission (11–13). Together, these observations point to a virus that is both ecologically widespread and highly promiscuous in its host associations. Despite this, JCV remains comparatively understudied, even as expanding vector ranges and documented reassortment among orthobunyaviruses raise concern for the emergence of novel variants (14–18).

Underlying this broad host range is a compact, tri-segmented RNA genome that supports replication in both mosquito vectors and mammalian hosts. JCV possesses a single-stranded, negative-sense RNA genome consisting of the small (S), medium (M), and large (L) segments. The S segment encodes the nucleocapsid (N) protein and the nonstructural protein NSs (19). Based largely on studies of La Crosse and Bunyamwera viruses, NSs is believed to modulate host immune responses to promote infection in both mammals and mosquitoes (20, 21). The L segment encodes the viral RNA-dependent RNA polymerase. The M segment encodes a glycoprotein precursor that is co-translationally processed into the surface glycoproteins Gn and Gc, as well as the nonstructural protein NSm. Gn and Gc assemble as heterodimers that form the viral envelope spikes and Gc is classified as a class II fusion protein (19). Neutralizing antibody responses are directed primarily against epitopes within the surface-exposed head domain of Gc (22). NSm is the least well-characterized of these proteins, and its function appears to be highly dependent upon the infected host with proposed roles ranging from assembly and morphogenesis, to transmission and dissemination (23–25). Together, proteins encoded on the S and M segments mediate viral entry, membrane fusion, assembly, intracellular trafficking and innate immune evasion, and therefore directly interface with host cell machinery. As such, they are likely primary determinants of viral fitness across both mosquito and mammalian hosts.

To investigate how viral genetics influence replication in mosquito and mammalian systems, we compared two distinct strains of JCV. JCV/61 is the prototypic 1961 isolate from Jamestown, Colorado, whereas JCV/03 is a more contemporary strain isolated in 2003 from pooled mosquitoes in Connecticut (26). Initial characterization by Bennett et al. (27) reported the strains share 91%, 83%, and 85% nucleotide identity across the small, medium, and large segments, respectively, with amino acid identity ranging from 91–98% depending on the protein. Despite this high degree of genetic similarity, JCV/61 and JCV/03 have been reported to exhibit distinct phenotypes in weanling Swiss Webster mice (27). This divergence suggests that relatively limited genetic differences underlie measurable phenotypic variation, making this strain pair a useful resource for identifying viral determinants of host-specific replication.

To identify JCV genetic determinants of host adaptation, we compared these two genetically similar strains of JCV. From this comparison we observed that the growth of JCV/61 and JCV/03 in mammalian Vero cells and neuronal cells was similar. In contrast, in mosquito-derived *Aedes albopictus* C6/36 cells we observed a striking lag in replication followed by a rapid increase (burst) in growth that was unique to the JCV/03 strain and dependent upon the cellular density. Furthermore, we found that a single passage in C6/36 cells conferred a heritable enhancement of JCV/03 replication in mosquito cells and eliminated the lag in growth. Illumina next-generation sequencing of mosquito cell-adapted JCV/03 identified a small number of candidate mutations in the M segment. Finally, we used a plasmid-based reverse genetics system to generate recombinant viruses carrying individual M segment mutations and directly test their contributions to viral replication in insect cells. Together, these results identify specific mutations in the M segment that modulate replication efficiency in mosquito cells and reveal multiple genetic pathways by which JCV can adapt to its vector host.

## 2. MATERIALS AND METHODS

### 2.1. Cells and viruses

Vero E6 cells (ATCC CRL-1586) and BSR-T7/5 cells were maintained in Dulbecco’s modified Eagle medium (DMEM) (Corning, Corning, NY, USA) supplemented with 10% cosmic calf serum (CCS) (HyClone, Cytiva, Marlborough, MA, USA) at 37 °C with 5% CO₂. BSR-T7/5 cells were supplied by Carolina Lopez (Washington University School of Medicine, St. Louis) and maintained under selection with 2 mg/mL Geneticin (Invitrogen, Eugene, OR, USA), added every two or three passages. C6/36 cells (*Aedes albopictus*) were acquired from Sara Cherry (University of Pennsylvania) and cultured in Leibovitz’s L-15 medium (Gibco, Waltham, MA, USA) supplemented with 10% fetal bovine serum (FBS) (Corning), 1% penicillin-streptomycin (Gibco), and 1% GlutaMAX (Gibco) and incubated at 28 °C with 5% CO₂. Primary rat cortical neurons were isolated from embryonic day 18 (E18) Sprague Dawley IGS rat pups (Charles River) by the Penn Medicine Translational Neuroscience Core as previously described (28). Neurons were maintained in Neurobasal medium (Gibco) supplemented with 2% B-27 (Invitrogen), 33 mM glucose, 40 mM HEPES (Gibco), and 1% penicillin-streptomycin and cultured in humidified chambers at 37 °C. Cultures were maintained for 7 days prior to use, with 40% of the medium replaced on day 4. All neuron media was conditioned at 37 °C with 5% CO₂ prior to use.

Jamestown Canyon virus (JCV) strain 61V-2235 (NR-536) was obtained from the NIH Biodefense and Emerging Infections Research Resources Repository, and JCV/03/CT was provided by Philip Armstrong (The Connecticut Agricultural Experiment Station). Both viral strains were purified by three rounds of terminal dilution as previously described (27). Four biological clones of each virus were generated and sequenced as described below. Working virus stocks were propagated in Vero E6 cells and titrated by quantitative PCR (qPCR), or plaque assay on Vero E6 cells, as described below.

### 2.2. Sequencing of purified viral clones

Four independently isolated biological clones of JCV/61 and JCV/03 were sequenced using a combination of cloning and PCR. Briefly, viral RNA was isolated from supernatant collected from infected cells using the Qiagen RNeasy Mini Kit (Cat #74104) and Buffer RLT Plus (Qiagen, Cat #1053393). cDNA was generated using the SuperScript™ III First-Strand Synthesis System (Invitrogen) and segment specific primers. The same segment-specific primers used to synthesize the cDNA were also employed to generate the PCR product. A complete list of these primers is documented in Supplemental Table 1.

To sequence the small segment, cDNA was amplified by PCR, then cloned into the pCRII-Blunt-TOPO plasmid using the Zero Blunt™ TOPO™ PCR Cloning Kit (Invitrogen, Cat #450245) according to the manufacturer’s protocol. To sequence the medium and large segment, cDNA was amplified by PCR and the linear amplicons were directly submitted for sequencing. Plasmids and linear amplicons were sequenced by Plasmidsaurus (Louisville, KY) using Oxford nanopore sequencing.

The generated consensus sequences were aligned to the existing reference genome using Geneious Prime (Version 2026.0.2). The clone with the fewest mismatches to the reference sequence was chosen as our prototype strain of JCV/61 and JCV/03. The reference for the small and medium segment of JCV/61 came directly from the data sheet supplied by BEI. The reference sequence for the large segment of JCV/61 and all three segments of JCV/03 were found in the literature (27). Supplemental Table 2 summarizes differences between the reference and the chosen prototype strain.

### 2.3. Plaque Assay

Infectious units were determined by plaque assay on confluent Vero E6 cells in 6-well plates. Virus stocks were serially diluted 10-fold in serum-free DMEM, and 850 µL of each dilution was added per well. Virus was adsorbed for 1 hour at 37 °C, after which the virus-containing medium was removed, rinsed once with Dulbecco’s phosphate-buffered saline (DPBS) (Corning), and cells were overlaid with a 1:3 mixture of 2.5% Avicel RC-591 (DuPont, Wilmington, DE, USA) and complete DMEM. Plates were incubated at 37 °C for 5 days. 4% paraformaldehyde was added directly to each well to fix the cells. After 1 hour the Avicel-paraformaldehyde overlay was removed and cells were stained with 0.1% crystal violet (Sigma-Aldrich, Burlington, MA, USA) in 20% methanol to visualize plaques. Plaques were manually counted, and counts from duplicate wells were averaged to calculate viral titers, which are reported as plaque-forming units per milliliter (PFU/mL).

### 2.4. Multistep growth curves in Vero, C6/36, and primary rat cortical neurons

Viral growth kinetics were assessed using multi-step growth curves in three cell types. C6/36, Vero, and primary rat cortical neurons were seeded in 6-well plates at 1.575 × 10^5^, 1.15 × 10^5^, and 4.8 × 10^5^ cells/mL, respectively. Neurons were grown on plates coated with 0.25 mg/mL poly-L-lysine hydrobromide (Sigma-Aldrich). One day after plating, cells were infected at multiplicity of infection (MOI) 0.1 or 0.03 in 2 mL complete, cell-type specific media. The virus-media inoculum was not removed, and titers were reported as fold-change relative to zero hour post-infection. At each timepoint, a small sample of supernatant was collected and replaced with fresh, cell-type specific media. Viral titers were quantified by plaque assay on Vero cells.

### 2.5. Analysis of replication in C6/36 Cells

To analyze JCV replication in C6/36 cells, infections were performed in 6-well plates with an initial plating density of 9 × 10⁵ cells/mL unless otherwise specified. Cells were incubated overnight at 28°C with 5% CO₂ and then infected with 2 × 10⁶ genome copies in 2 mL of complete medium per well. C6/36 cells were minimally attached and prone to lifting, and due to the importance of cell density to viral growth kinetics, virus inoculum was not removed following infection. After infection, plates were returned to the incubator for the duration of the time course. An aliquot of the diluted virus (approximately 1 × 10⁶ genome copies/mL) was retained as the zero-hour input sample and used for fold-change calculations.

At each indicated time point, 75 µL of culture supernatant was collected and clarified by centrifugation at 1,200 × g for 5 min. An equal volume of fresh medium was added back to each well. Following clarification, 50 µL of supernatant was diluted into DPBS to a final volume of 140 µL and stored at −80°C until RNA extraction.

For matched extracellular and intracellular time course experiments, the entire volume of cell supernatant was removed at each time point and clarified by centrifugation at 1,200 × g for 5 min. Following supernatant removal, cells were lysed directly in the culture wells by addition of 1 mL of TRIzol ™ Reagent (Invitrogen). Clarified supernatant and cell lysates were stored at −80°C until processing.

For variable-density growth curve experiments, cells were plated at high, medium, or low densities and infected with an equal number of plaque forming units, resulting in MOIs of 0.03, 0.01, and 0.003, respectively. The supernatant was sampled over time as described above.

### 2.6. Analysis of replication in Vero Cells

To analyze replication in Vero E6 cells, cells were plated at a density of 1 × 10⁵ cells/mL in complete DMEM and incubated overnight at 37°C with 5% CO₂. Cells were infected with 2 × 10⁶ genome copies in 2 mL of serum-free DMEM and incubated at 37°C for 1 hour to allow virus entry. With the exception of serial passaging experiments, the virus inoculum was removed, cells were gently rinsed once with DPBS, and 2 mL of complete medium was added to each well, then the plates were returned to the incubator for the duration of the time course. At each indicated time point, 75 µL of culture supernatant was collected, clarified by centrifugation at 1,200 × g for 5 min, and replaced with an equal volume of fresh complete medium. Clarified supernatants were stored at −80°C until RNA extraction.

### 2.7. RNA Extraction and cDNA synthesis

Viral RNA was extracted from clarified supernatants using the QIAamp Viral RNA Mini Kit (Qiagen, Cat #52906) according to the manufacturer’s instructions and eluted in 30 µL of supplied AVE buffer. Total cellular RNA was isolated from TRIzol ™ Reagent (Invitrogen) lysates and further purified using the Zymo RNA Clean & Concentrator Kit (Zymo Research Cat # R1018, Irvine, CA, USA), with on-column DNase I treatment (Zymo Research, Cat #E1010). Purified cellular RNA was quantified using the Qubit RNA High Sensitivity Assay Kit (Invitrogen) on a Qubit 4 Fluorometer (Invitrogen).

For cDNA synthesis, either 8 µL of viral RNA from supernatants or equal amounts of total cellular RNA were reverse transcribed using the SuperScript™ III First-Strand Synthesis System (Invitrogen) with random primers (Invitrogen, Cat #4819011), following the manufacturer’s instructions.

### 2.8. Construction of JCV/03-pN plasmid for qPCR standard curve

The JCV-pN expression plasmid used to generate the qPCR standard curve for absolute genome copy number quantification contains the JCV/03 nucleocapsid gene cloned into the pIRES-polyA backbone (constructed by Paul Bates, University of Pennsylvania). The pIRES-polyA plasmid backbone and JCV/03 nucleocapsid open reading frame were amplified using Q5 High-Fidelity 2X Hot Start Master Mix (New England Biolabs Cat #M0494S, Ipswich, MA, USA) and the primers listed in Supplemental Table 3. Primer overhangs were designed to generate regions of homology required for HiFi DNA Assembly (NEB, Cat #M5520AA). The JCV/03 nucleocapsid PCR template was the gBlock™ (Integrated DNA Technologies, Coralville, IA, USA) used to construct the JCV/03-pS launch plasmid. The linearized backbone and PCR-amplified insert were assembled using NEBuilder HiFi DNA Assembly (NEB) according to the manufacturer’s instructions.

### 2.9. qPCR

Viral genome copy number in supernatant and cellular RNA was quantified by qPCR using PowerUp™ SYBR™ Green Master Mix (Applied Biosystems, Waltham, MA, USA) and JCV-specific primers. Primers targeted the small genome segment and were designed to bind identical regions in both strains to ensure comparable amplification efficiency. Supplemental Table 4 lists the qPCR primers used in this study.

Absolute genome copy number was quantified using a six-point standard curve generated from 10-fold serial dilutions of the JCV/03-pN expression plasmid. Plasmid concentration was converted to copy number using a molecular weight-based equation (30). qPCR was performed on a QuantStudio 6 Flex Real-Time PCR System (Applied Biosystems) using the following cycling conditions: initial incubation at 50°C for 2 min and 95°C for 10 min, followed by 40 cycles of 95°C for 15 sec, 52°C for 30 sec, and 72°C for 30 sec, with fluorescence acquisition during the 52°C and 72°C steps.

For samples derived from cell lysates, values were normalized to actin, using a ΔCt-based correction (Ct_actin – Ct_ref, where Ct_ref is the time-matched mock), yielding housekeeping-adjusted genome copies per µg RNA. For samples derived from culture supernatants, normalization to a housekeeping gene was not possible; therefore, viral genome copies per milliliter were calculated based on the volume of supernatant collected and the volume of cDNA input, assuming uniform recovery and reaction efficiency across samples and processing steps.

### 2.10. RNA sequencing and alignment

Viral RNA was extracted from clarified cell culture supernatants using TRIzol™ LS Reagent (Invitrogen) and purified using the Zymo RNA Clean & Concentrator kit (Zymo Research). Purified RNA was submitted to the University of Pennsylvania School of Veterinary Medicine Center for Host–Microbial Interactions for library preparation and sequencing. cDNA libraries were prepared using the Illumina® Stranded Total RNA Prep Ligation with Ribo-Zero Plus kit (Illumina Cat #20040529, San Diego, CA, USA) and sequenced on an Illumina NextSeq 2000 platform using a P2 100-cycle kit (Illumina, Cat #20046811).

Raw sequencing reads were aligned to the published Jamestown Canyon virus reference genome (GenBank accession number by segment, Small: HM007353.1, Medium: HM007354.1, Large: HM007355.1) using breseq (v0.36.0) in polymorphism mode, which assumes a mixed population of viral variants. Read mapping was performed using Bowtie2, and mutations were identified relative to the reference genome using a minimum variant frequency cutoff of 5%. The breseq gdtools compare function was used to generate comparative mutation frequency reports across samples (31).

### 2.11. Construction of JCV/03 launch plasmids

Launch plasmids encoding the JCV small, medium, and large genome segments were constructed using a modular DNA assembly approach. A T7-driven plasmid backbone containing the T7 optimal promoter, super cut hepatitis delta virus (HDV) ribozyme, and T7 terminator was used to clone the S, M and L segment inserts. Viral genome sequences corresponding to JCV/03 (Small: HM007353.1, Medium: HM007354.1, Large: HM007355.1) were synthesized as double-stranded DNA fragments (gBlocks™, Integrated DNA Technologies) containing the antigenomic sequence of each genome segment with the addition of the hammerhead ribozyme sequence positioned at the 5′ end. The complete S segment was synthesized as a single fragment, the M segment as two fragments, and the L segment as four fragments. All fragments were designed with overlapping regions compatible with seamless assembly. Plasmids pS, pM, and pL were assembled by combining the linearized plasmid backbone with the corresponding PCR-amplified gBlocks™ using NEBuilder HiFi DNA Assembly (NEB) according to the manufacturer’s instructions. Final constructs were verified by Oxford Nanopore (Eurofins Genomics, Louisville, KY)

### 2.12. Construction of M segment mutants

Individual point mutations within the pM plasmid were introduced by PCR-based mutagenesis using primers listed in Supplemental Table 5, followed by circularization using NEBuilder HiFi DNA Assembly (NEB). All PCR reactions described in this manuscript were performed using Q5 High-Fidelity 2X Hot Start Master Mix (NEB) according to the manufacturer’s instructions.

### 2.13. Rescue of recombinant virus

Rescue of recombinant virus was performed in BHK BSRT7/5 cells using Lipofectamine L3000 (Invitrogen, Cat #L3000015). Cells were seeded at a density of 1.8 × 10⁵ cells/mL and transfected with equal amounts (0.5-2 ug each) of plasmids JCV/03-pS,-pM, and-pL. Transfections were carried out according to the manufacturer’s instructions, using 1 µL Lipofectamine 3000 and 1 µL P3000 per µg of total DNA. 3-5 days post-transfection, cells and supernatants were pooled with approximately 1 × 10⁵ Vero cells, plated with fresh growth medium in a 10 cm dish, and returned to the incubator. When roughly 70% cytopathic effect was observed, cultures were harvested, clarified, and aliquoted as passage 0 (P0) virus stocks. Passage 1 (P1) stocks were generated by amplification on Vero cells and experiments were performed with P1 viral stocks.

To verify successful rescue of recombinant viruses containing point mutations, viral RNA was extracted from P0 and P1 stocks and used for cDNA synthesis as described above. cDNA was used as template for PCR targeting regions within the M segment containing the mutation of interest, using the primers in Supplemental Table 1. Linear amplicons were submitted to EuroFins (Eurofins Genomics, Louisville, KY) for Oxford nanopore sequencing.

## 3. RESULTS

### 3.1. JCV/03 displays a mosquito cell-specific replication delay

JCV/03 and JCV/61 had previously been reported to have differing *in vivo* pathology phenotypes with JCV/61 being more neuropathogenic, however they were reported to be similar when analyzed *in vitro* (27). To begin our evaluation of these viruses, we performed limited dilution cloning on Vero E6 cells to produce uniform viral stocks. One clone for each virus strain was then chosen for analysis of growth *in vitro* in three cell lines. African green monkey Vero E6 cells, mosquito-derived *Aedes albopictus* C6/36 cells, and primary rat cortical neurons were infected with JCV/61 or JCV/03 at multiplicity of infection (MOI) 0.1 and 0.03, and released virus was sampled at regular intervals for the duration of cell viability. Viral titer at each timepoint was quantified by plaque assay on Vero E6 cells. In Vero cells JCV/03 and JCV/61 exhibited similar replication kinetics. The replication curve profiles were comparable at both MOIs, with JCV/03 ultimately reaching a higher peak titer than JCV/61 (Figure 1A). In primary rat cortical neurons both viruses initially replicated with similar kinetics, reaching a peak titer at day 2. As seen with replication in Vero cells, JCV/03 titers were higher than those of JCV/61. Additionally, JCV/61 titers declined at day 3 due to neuronal cell death, while JCV/03 titers remained relatively constant for the duration of the time course through day 7, at which point the neuronal cells began losing viability (Figure 1B).

**Figure 1:**
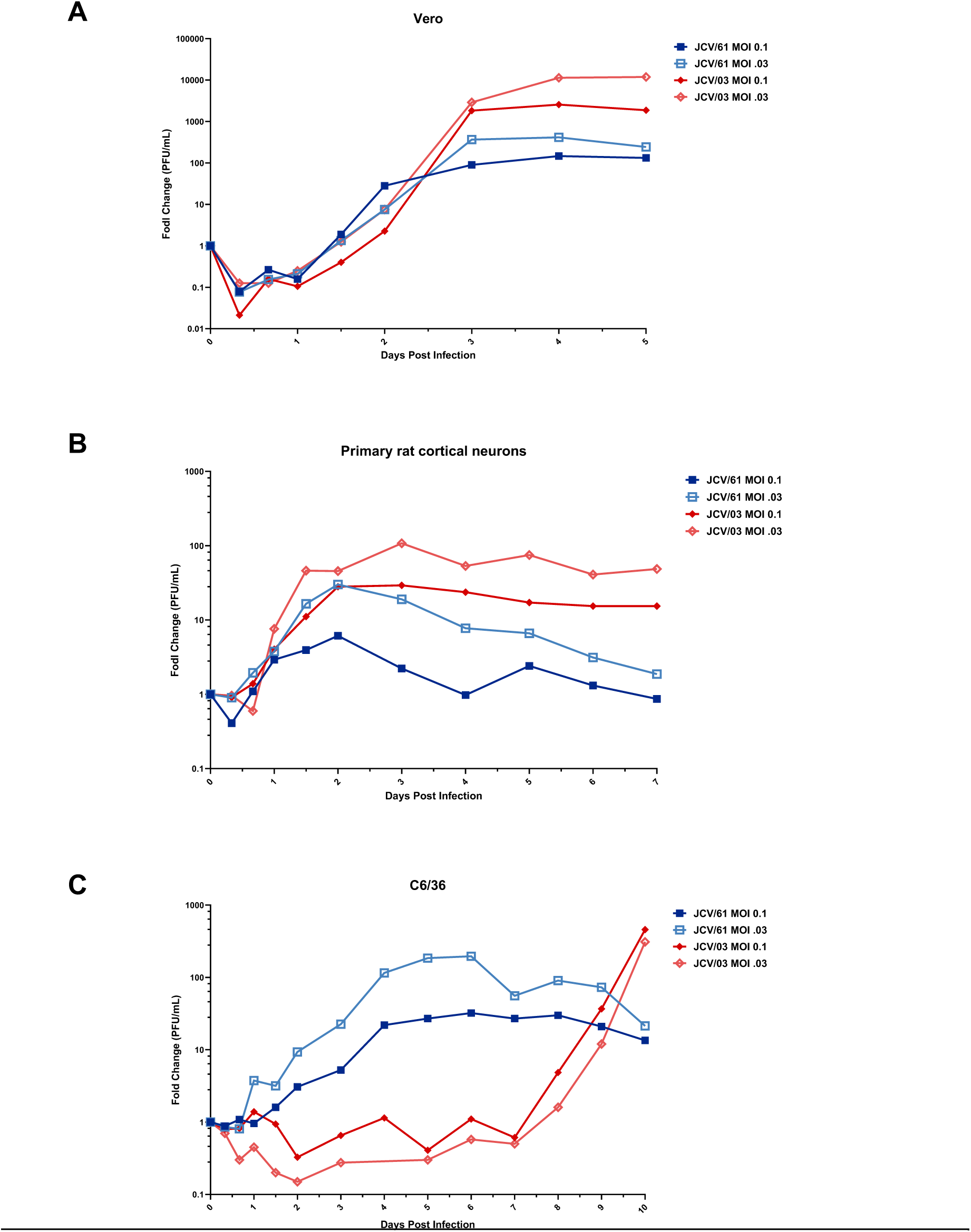
JCV/03 exhibits a distinct lag-burst growth phenotype in C6/36 cells. A) Vero cells, B) primary rat cortical neurons, and C) C6/36 cells were infected with two strains of JCV at MOI 0.1 and MOI 0.03. Supernatant was collected at indicated time points and infectious virus was quantified by plaque assay on Vero cells. Titers are expressed as fold change in PFU/mL relative to input.

In contrast to what is observed in mammalian cells, replication in mosquito C6/36 cells revealed a striking strain-specific difference between JCV/61 and JCV/03 (Figure 1C). JCV/61 replicated steadily, reaching peak titers by approximately 4 days post-infection and subsequently plateauing. JCV/03 however, exhibited minimal accumulation of infectious virus for seven days post-infection, followed by an abrupt transition to rapid replication. During the final three days of the time course, JCV/03 titers increased nearly 1000-fold, ultimately surpassing JCV/61. This “lag-burst” pattern was identical between the two MOIs. Given this unusual behavior, we analyzed replication in mosquito cells for three additional purified clones of JCV/03 and the uncloned parental stock that was obtained from The Connecticut Agricultural Experiment Station. All four clones and the parental stock displayed a six to eight day delay in replication followed by a dramatic replication increase (Supplemental Figure 1). Thus, the unusual phenotype for JCV/03 replication in mosquito cells is an inherent feature and not restricted to a selected clone. Overall, these results establish that while JCV/03 and JCV/61 replicate similarly in mammalian cells, JCV/03 exhibits a distinct lag-burst phenotype exclusively in insect C6/36 cells.

### 3.2. The JCV/03 lag-burst phenotype reflects delayed viral RNA replication

Analysis of JCV/03 virus in the supernatant of infected insect cells revealed an unusual delay in appearance of infectious virus, therefore we wanted to address whether JCV/03 was replicating intracellularly but failing to efficiently assemble or exit cells. C6/36 cells were infected with equal genome copies of JCV/61 or JCV/03, and at each time point the entire supernatant and corresponding cell lysates were collected for quantification of viral RNA. As observed previously for infectious virus, JCV/61 genomic RNA in the supernatant increased steadily over time, whereas the release of JCV/03 genomic RNA was delayed until day 8 when it rapidly increased (Figure 2A, compare blue vs red bars). qPCR analysis of intracellular RNA revealed patterns similar to that of released virus. JCV/61 genomic RNA reached peak levels early on and remained stable for the duration of the time course (Figure 2B, blue bars). As was seen for extracellular virus, JCV/03 showed minimal intracellular viral RNA accumulation during the first 6 days, followed by a rapid increase at day 8 to a peak that was comparable to JCV/61, which then plateaued (Figure 2B, red bars). Therefore, the delayed kinetics of JCV/03 were evident both intracellularly and extracellularly, indicating the lag phase reflects delayed viral replication rather than a lag in viral assembly or egress.

**Figure 2:**
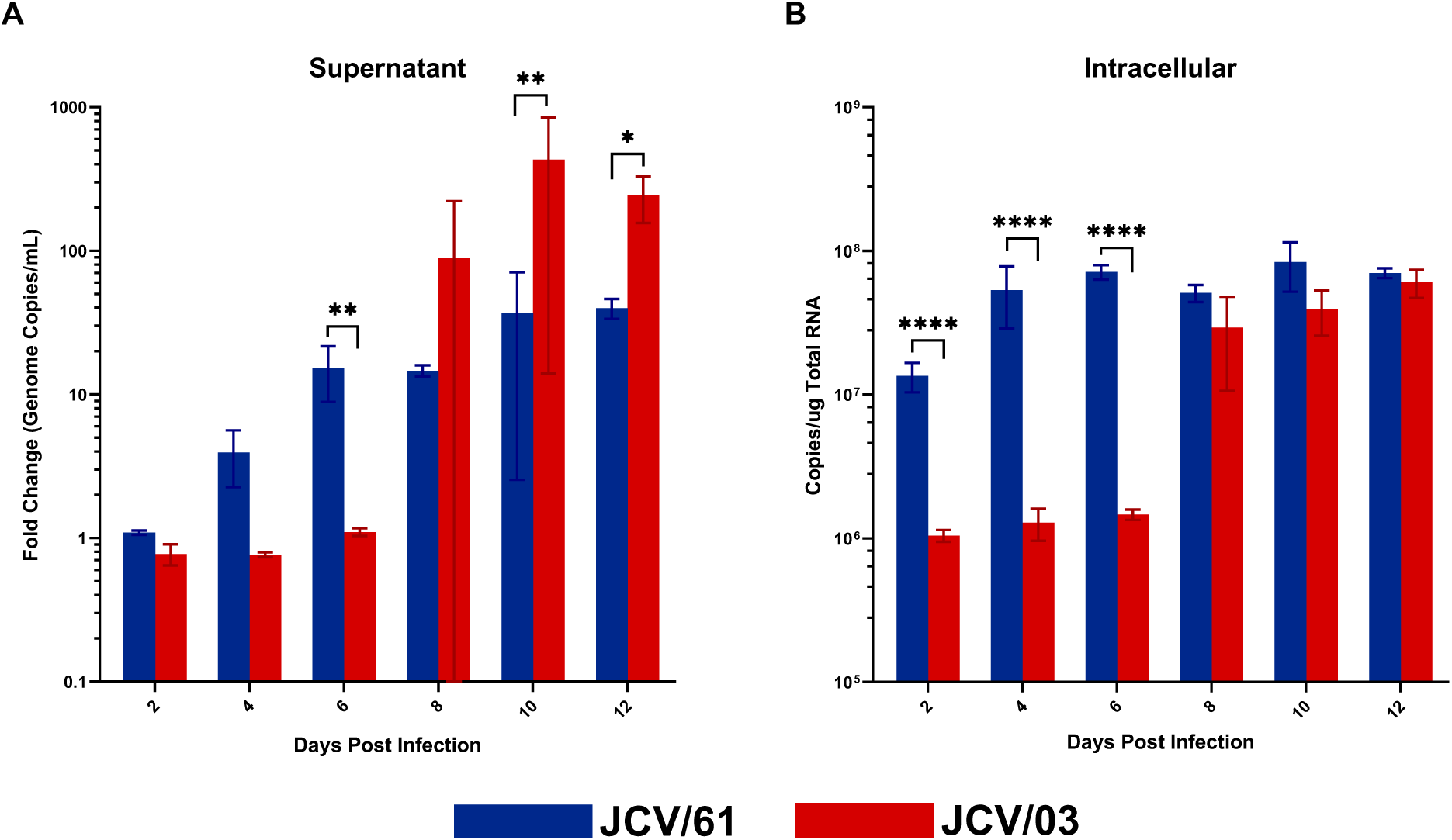
The JCV/03 lag–burst phenotype reflects delayed viral replication. C6/36 cells were infected with 2 × 10⁶ genome copies of JCV/61 or JCV/03. (A) Extracellular viral RNA was quantified from collected supernatants by qPCR and expressed as fold change in genome copies per mL relative to input. (B) Intracellular viral RNA was quantified from total cellular RNA at corresponding time points and expressed as genome copies per μg total RNA. Data represent mean and SD of three biological replicates. Statistical significance between viruses at each time point was determined by ordinary two-way ANOVA with Sidak’s multiple comparisons test performed on log10-transformed values. *P < 0.05, **P < 0.01, ***P < 0.001.

### 3.3. Plating density modulates timing and magnitude of JCV/03 replication in C6/36 cells

The abrupt transition of JCV/03 to rapid replication suggested that an extrinsic, time-dependent factor might govern viral growth, such as cell-cell proximity, abundance or accumulation of secreted factors that affect cellular physiology. If this were the case, then we hypothesized that increasing the initial cell density might accelerate the transition to rapid viral replication. To address whether cellular density affected JCV/03 replication, C6/36 cells were plated at low, medium, or high density and infected with a fixed quantity of infectious JCV/61 or JCV/03. JCV/61 replication was consistent across conditions and growth curves segregated according to initial plating density, with higher-density wells yielding higher overall genome accumulation but no qualitative change in replication kinetics (Figure 3, blue curves). By contrast, JCV/03 replication was strongly influenced by plating density. Increasing cell density progressively shortened the lag phase, however after the lag, the slope of the curves appeared similar. The plating density also appeared to alter the magnitude of replication. The highest-density condition exhibited the earliest transition to exponential growth at day 6 and reached the highest peak viral titer. Compared to JCV/61 plated at the same cell density, this was an approximately 100-fold higher peak titer. (Figure 3, dark red vs dark blue curves). These data demonstrate that cell density is directly correlated with both the timing and magnitude of the JCV/03 lag-burst phenotype, although the mechanistic basis of this relationship remains undefined.

**Figure 3:**
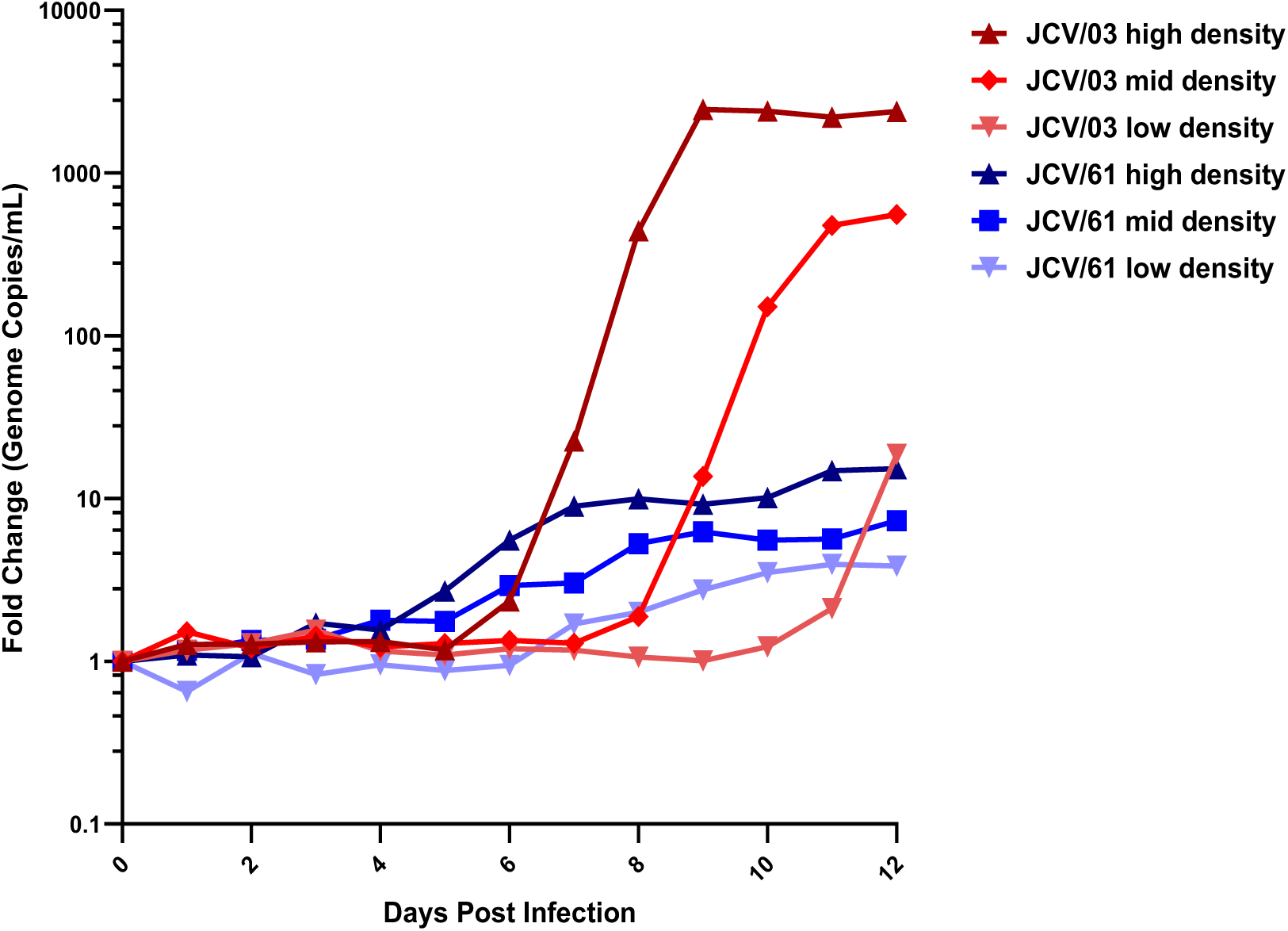
Cell density modulates JCV/03 replication kinetics in C6/36 cells. C6/36 cells were seeded in 6-well plates at low (2 × 10⁵), mid (6 × 10⁵), or high (1.8 × 10⁶) density (cells per well) and infected with 6,000 PFU of JCV/61 or JCV/03. The resulting MOIs were 0.03, 0.01, and 0.003 for low-, mid-, and highdensity wells, respectively. Supernatants were collected at the indicated time points, and viral titers were quantified by qPCR to determine genome copies per.

To further investigate extracellular factors governing the lag-burst replication phenotype, infections were performed in C6/36 cells cultured in buffered or conditioned media. Buffered media consisted of standard C6/36 growth media supplemented with HEPES to a final concentration of 25 mM. The conditioned media was developed by plating C6/36 cells at high density and collecting the media 12 days post mock infection. Three independent batches of conditioned media were tested. In standard media, JCV/03 exhibited minimal replication through day 6 post infection, rapidly reached peak titers by approximately day 8, and then remained stable through day 12 (Figure 4, red line). In buffered media, the lag phase decreased modestly, with titers beginning to increase by day 4 post infection; however, the virus still reached peak titers on day 8 and to the same magnitude as in standard media (Figure 4, blue line). In conditioned media batches A and B, the replication lag was similarly shortened, with titers increasing by day 4 and plateauing by day 8. Due to cytotoxicity induced by the conditioned media, peak titers were approximately ten-fold lower than those observed in standard or buffered media (Figure 4, green and orange lines). In contrast, conditioned media batch C displayed no lag in replication, but the pace was gradual rather than exponential, with titers steadily increasing through day 6 before plateauing (Figure 4, purple line). Together, these findings indicate that while extracellular conditions can influence the timing and magnitude of JCV/03 replication in C6/36 cells, they are insufficient to eliminate the lag-burst pattern, suggesting that this replication phenotype is primarily driven by intrinsic viral determinants.

**Figure 4:**
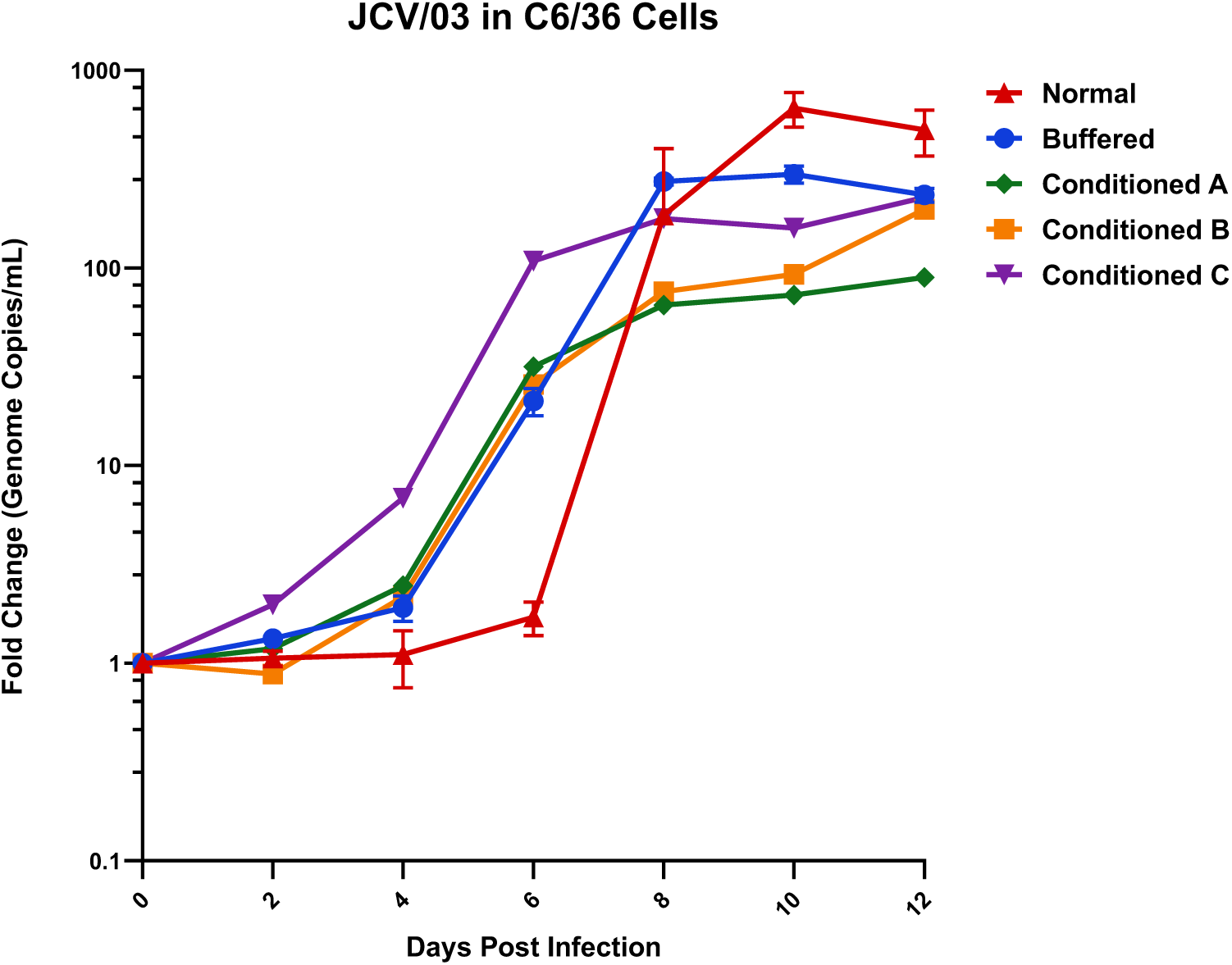
Altering media conditions modulates JCV/03 replication kinetics in C6/36 cells. C6/36 cells were infected with 2 × 10⁶ genome copies of JCV/03 in standard growth media, growth media buffered with 25 mM HEPES, or conditioned media collected from high-density mock-infected C6/36 cultures. Three batches of conditioned media were tested. Viral genome copies in culture supernatants were quantified over time by qPCR and expressed as fold change relative to input. One biological replicate per batch of conditioned media. Two biological replicates of normal and buffered media.

### 3.4. A single passage in C6/36 cells confers heritable enhancement of JCV/03 replication

We next asked whether the abrupt increase in JCV/03 replication in C6/36 cells reflected a heritable change and whether this adaptation depended on the initial cellular plating density. To address this, parental stocks of JCV/03 and JCV/61 were used to infect C6/36 cells plated at high-and low-density and virus was collected from these cultures on day 12. Equal genome copies of these mosquito cell grown (passaged) virus stocks were then used to infect fresh C6/36 cells plated at high density. For comparison, C6/36 cells were also infected with equal genome copies of the parental JCV/61 or JCV/03 stocks that were derived from Vero cells. Mosquito cell-derived JCV/61 virus exhibited a modest enhancement of replication when used to infect fresh C6/36 cells (Figure 5, dashed blue lines). Compared to Vero-derived virus, replication of JCV/61 from mosquito cells was slightly accelerated and peak titers at day 12 were approximately 5-fold higher (Figure 5, compare solid and dashed blue lines), but overall the replication pattern was unchanged. In contrast, JCV/03 collected from both high-and low-density C6/36 cultures replicated immediately and exponentially when infecting fresh C6/36 cells, with no detectable delay (Figure 5, dashed red lines). Peak titers were reached by day 6 and leveled off. As seen before, Vero-derived JCV/03 displayed a distinct replication lag, transitioned to rapid replication on day 6, and then plateaued through the end of the 12 day experiment (Figure 5, solid red line). Across numerous independent experiments, we consistently observed that a single passage in C6/36 cells was sufficient to enhance JCV/03 replication in mosquito cells. Collectively, these data demonstrate that JCV/03 acquires heritable, enhanced mosquito-cell replication after a single passage in C6/36 cells, regardless of the initial plating density.

**Figure 5:**
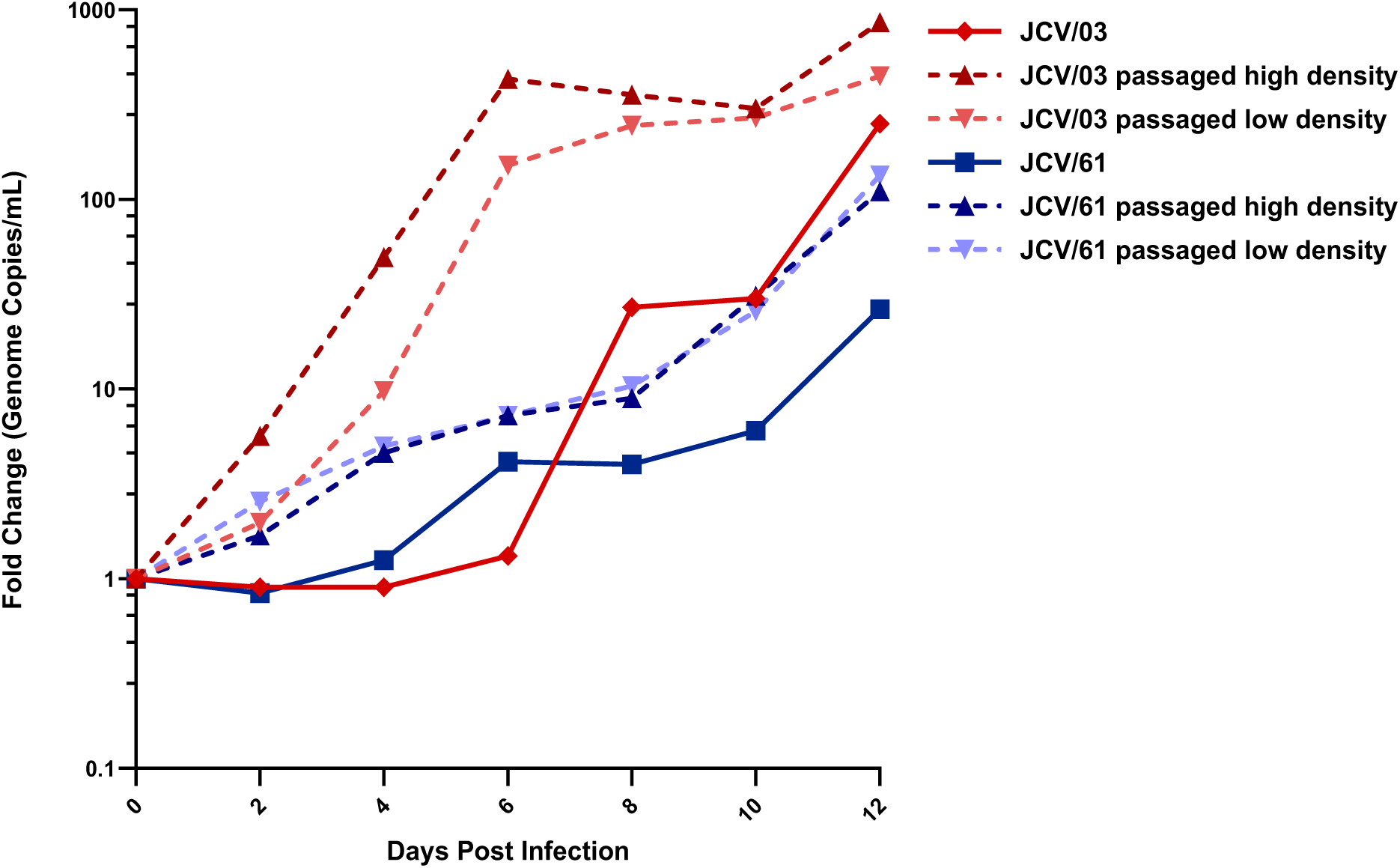
A single passage in C6/36 cells confers heritable enhancement of JCV/03 replication. C6/36 cells were infected with 2 × 10⁶ genome copies of parental JCV/03 or JCV/61, or with parental virus passaged once in C6/36 cells at low or high plating density. Supernatants were collected at the indicated time points, and viral RNA was quantified by qPCR to determine genome copies per mL. Titers are expressed as fold change in genome copies per mL relative to input.

### 3.5. Serial passage reveals distinct evolutionary trajectories of JCV/03 adaptation

Given that a single passage of JCV/03 in C6/36 cells conferred a heritable replication advantage in C6/36 cells, we next asked whether this adaptation affected viral replication in mammalian cells and whether the advantage would persist or revert following passage in mammalian cells. To address these questions, we performed two independent serial passage experiments and analyzed the viral phenotype and genotype after each passage. In each series, JCV/03 was passaged twice in high density C6/36 cells, followed by two passages in Vero cells, and a fifth passage in back into C6/36 cells (Figures 6A and 6B). The parental stock of Vero-derived JCV/03 was employed as the control for each set of infections (Figures 6A and 6B, solid red lines). During the first set of serial passages (Series I), Vero-derived JCV/03 displayed the characteristic replication lag followed by a rapid increase at day 8 (Figure 6A, first panel). As seen previously, mosquito cell-derived JCV/03 taken from passage 1 showed no replication lag when infecting fresh C6/36 cells (Figure 6A, second panel dashed line). Replication of JCV/03 in Vero cells during passages three and four was identical to parental Vero cell-derived virus (Figure 6A, dashed lines third and fourth panels). However, when the virus from Vero passage in panel 4 was returned to C6/36 cells, replication had reverted to the original lag-burst phenotype observed for parental JCV/03 (Figure 6A, panel five).

**Figure 6:**
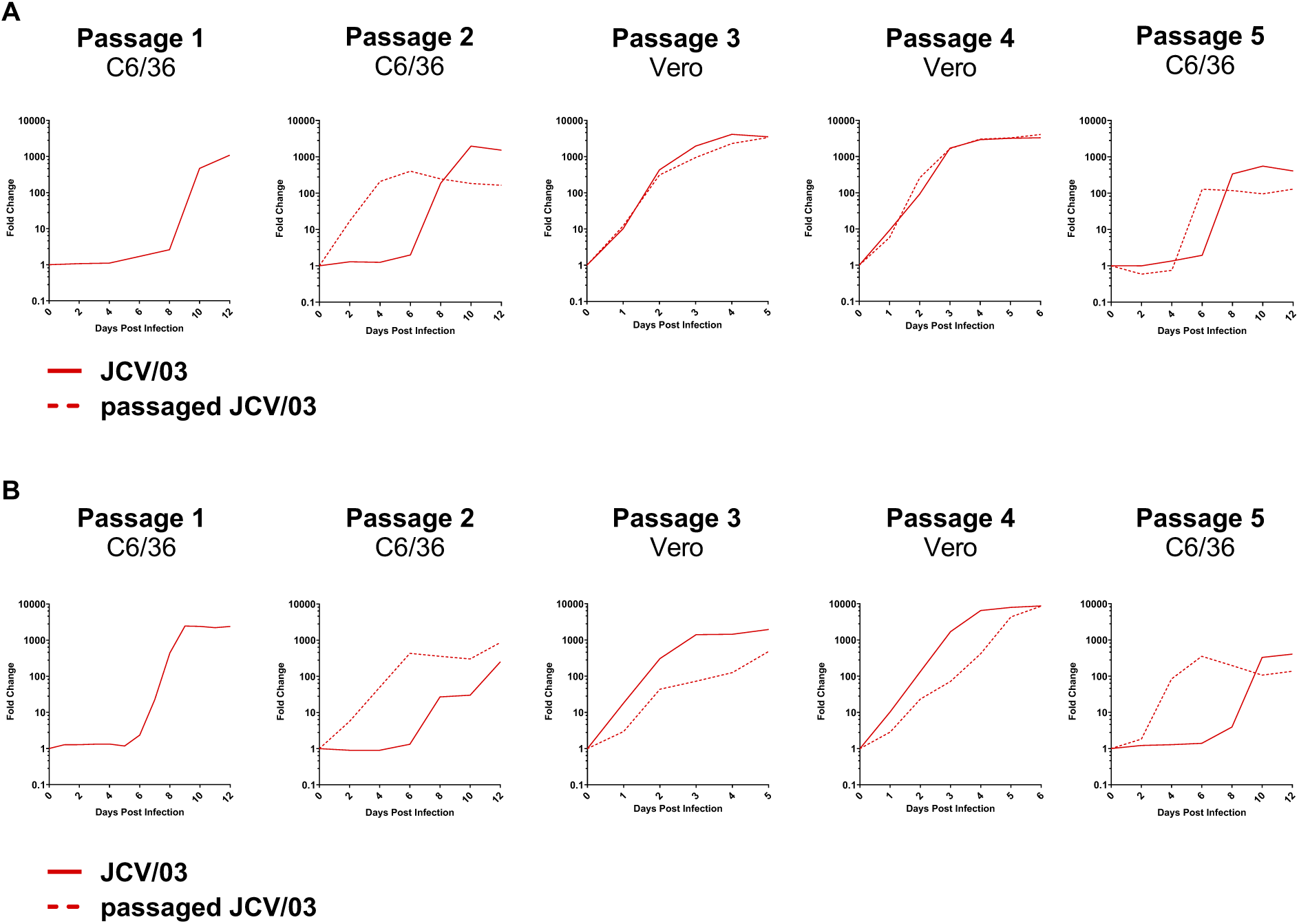
Serial passage reveals consistent adaptation in insect cells but divergent outcomes after mammalian passage. Two independent five-passage series were Performed. In each series, C6/36 or Vero cells were infected with 2 × 10⁶ genome copies of ither parental JCV/03 (solid line) or passaged JCV/03 (dashed line). Virus was passaged wice in C6/36 cells (Passages 1 and 2), followed by two passages in Vero cells (Passages 3 and 4), and subsequently returned to C6/36 cells (Passage 5). To generate growth curves for Series I (A) and Series II (B), supernatants were collected at he indicated time points and viral RNA was quantified by qPCR to determine genome copies er mL. Titers are expressed as fold change in genome copies per mL relative to input.

The second serial passage experiment (Series II) also showed viral adaptation to mosquito cells during passage 1 resulting in the absence of a lag phase when this stock was used to infect fresh C6/36 cells for passage 2 (Figure 6B, dashed line second panel). However, in contrast to the results from Series I, adaptation to C6/36 cells was accompanied by subsequent impaired replication in Vero cells, where mosquito cell passaged virus replicated slower than parental JCV/03 (Figure 6B, dashed lines third and fourth panels). Additionally, unlike what was observed in Series I, the Series II virus did not revert after two passages in Vero cells; when returned to C6/36 cells in passage five, replication was rapid and the lag phase was significantly abbreviated (Figure 6B, panel 5 dashed line). Overall, these results confirm that heritable changes occur in JCV/03 upon growth in mosquito cells and these changes can be stable (Series II) or revert (Series I) upon growth in mammalian cells.

To determine whether initial plating density influenced the outcome of serial passage, we repeated the five-passage experiment using virus initially adapted in low-density C6/36 cultures (Supplemental Figure 2). Passage 1 was performed with an initial plating density approximately six-fold lower than the high-density conditions used in Figure 6, while passages 2-5 were performed as described above. Similar to virus adapted using high-density mosquito cell cultures, low-density-derived JCV/03 displayed immediate replication upon reinfection into C6/36 cells, with no apparent lag phase (Supplemental Figure 2, panel 2). Although replication in passage 3 was comparable to that of parental Vero-derived virus, growth was noticeably reduced by passage 4, resembling the phenotype observed for the high-density virus in Series I. (Supplemental

Figure 2, panel 3 and panel 4). When returned to C6/36 cells for passage 5, the virus retained an abbreviated lag phase and accelerated replication kinetics, indicating that adaptation to insect cells persisted, at least partially, despite two passages in mammalian cells. This shortened lag is similar to that observed in the Series II passages. Together, these results demonstrate that the initial plating density does not substantially alter the long-term outcome of serial passage.

### 3.6. A plasmid-based reverse genetics system reveals S193G is required for the JCV/03 phenotype

To reveal changes responsible for adaptation to mosquito cells, JCV/03 viral RNA from the supernatant at the end of each passage (Figure 6, day 12 on insect cells and day 5 or 6 on Vero cells) was extracted and sequenced. Reads were aligned to the published JCV/03 reference genome (Small: HM007353.1, Medium: HM007354.1, Large: HM007355.1). During this analysis we identified a variation in the M segment region encoding Gn (S193G) that was fixed in our parental stock of JCV/03 and present in all four JCV/03 clones purified by limiting dilution but absent from the published reference genome (27). Upon growth in either Vero or mosquito cells, the S193G variant remained stable in both Series I and Series II. Because this change was present at 100% frequency in our parental JCV/03 stock and maintained at 100% throughout the serial passage experiments, we sought to determine if S193G contributed to the delayed replication phenotype observed in mosquito cells.

To investigate the importance of the changes identified in JCV/03, we employed a plasmid-based reverse genetics system to generate recombinant viruses with precisely defined genomes. In this system, T7-driven plasmids encoding each genome segment were co-transfected into BHK BSR-T7/5 cells, which constitutively express T7 RNA polymerase (32). The launch plasmids contained the full-length antigenomic cDNA of a genome segment, including the UTRs, positioned between an optimized T7 promoter and T7 terminator (33). To ensure precise termini of the viral RNA transcripts, plasmids were constructed with self-cleaving ribozymes flanking the genome segment. A hammerhead ribozyme was positioned upstream of the viral genome segment to generate the correct 5′ end, while a super cutter hepatitis delta virus (HDV) ribozyme was placed downstream to produce the correct 3′ end (34, 35). Following transfection, the T7 RNA polymerase expressed in BSR-T7/5 cells transcribed antigenomic cDNA from each plasmid, initiating translation of viral proteins and production of infectious virus. After launch, stocks of the recombinant viruses were produced by one round of replication in Vero cells.

Recombinant JCV/03 that encodes a serine at position 193 as indicated by the reference sequence (rJ.03-S) or a glycine as in our parental stock sequence (rJ.03-G) were launched and characterized. We compared our stock JCV/03, rJ.03-S, and rJ.03-G by plaque assay and multi-step growth curves in Vero and C6/36 cells. All three viruses produced mixed-size plaques on Vero cells (Figure 7A). Growth curves in both Vero and C6/36 cells revealed a significant difference between the two recombinant viruses. As might be expected, the rJ.03-G virus which is identical in sequence to the parental JCV/03 stock, replicated similarly to the parental virus in both mammalian (Figure 7B) and insect cells (Figure 7C). By contrast, rJ.03-S replicated more slowly in both insect and mammalian cells (orange curves, Figures 7B and 7C). Notably, in C6/36 cells rJ.03-S did not exhibit the characteristic lag-burst phenotype observed for parental JCV/03, while rJ.03-G displayed the parental phenotype (Figure 7C). Overall, these results demonstrate that a serine at position 193 is detrimental for replication in both mammalian and insect cells. Moreover, only virus with glycine in this position displays a replication lag in mosquito cells.

**Figure 7:**
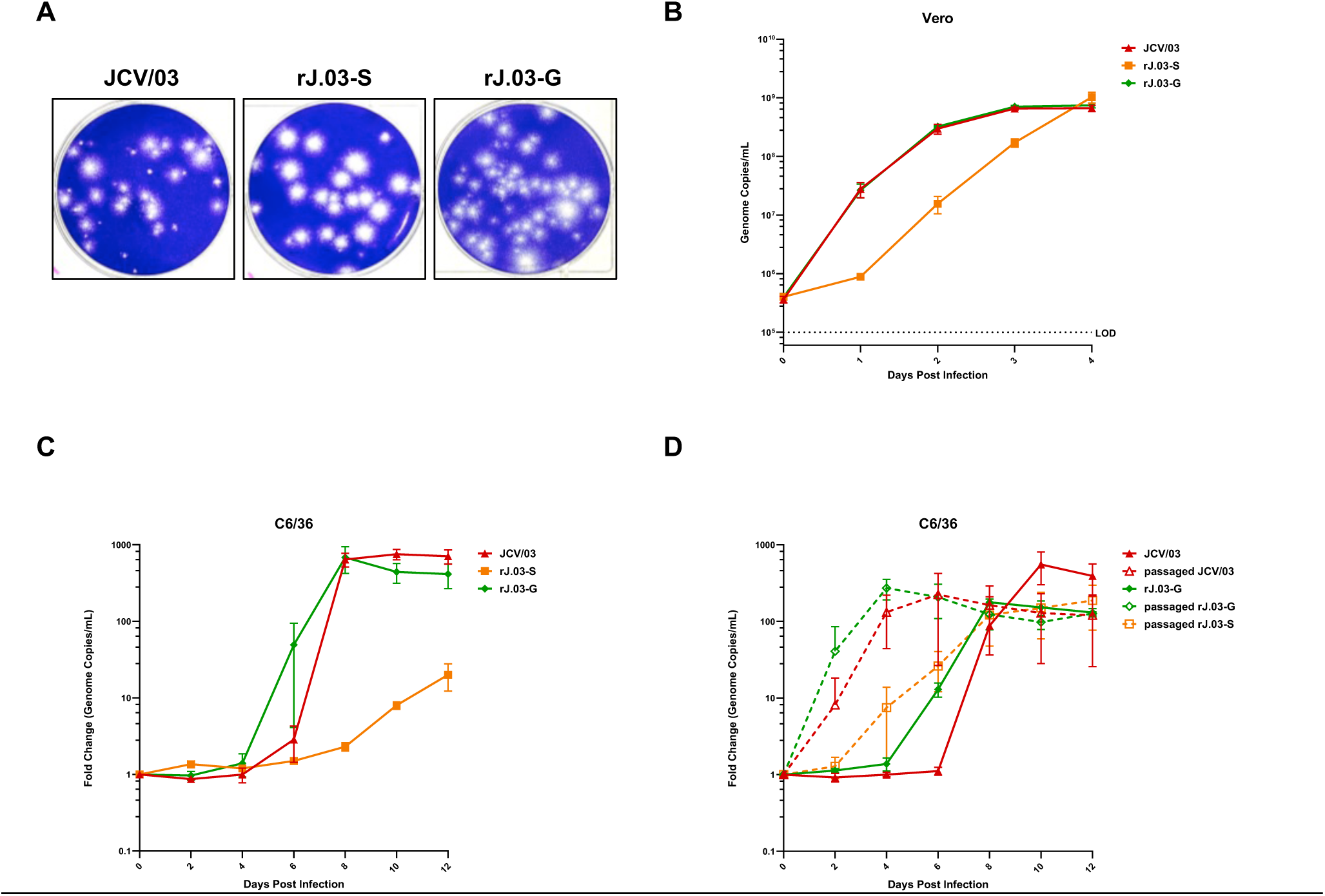
S193G is required for the parental JCV/03 lag–burst phenotype. (A) Representative plaque morphology of parental JCV/03, recombinant JCV/03 encoding serine at residue 193 (rJ.03-S), and recombinant JCV/03 encoding glycine at residue 193 (rJ.03-G) Following 5 days of growth on Vero E6 cells. Multi-step growth curves of parental JCV/03, J.03-S, and rJ.03-G in (B) Vero E6 cells and (C) C6/36 cells. (D) Growth of parental JCV/03 and unpassaged rJ.03-G compared with passaged JCV/03, passaged rJ.03-S, and passaged J.03-G in C6/36 cells. Supernatants collected at indicated timepoints; viral RNA quantified by qPCR to determine genome copies/mL. In C and D, titers expressed as fold change in genome copies/mL relative to input. Data represent mean and SD of three biological replicates. Dotted Line in B indicates limit of detection.

To determine whether S193G was also required for the heritable adaptation observed in C6/36 cells or only necessary for the initial lag-burst phenotype, we compared the growth of insect cell-derived rJ.03-S, rJ.03-G, and JCV/03 with Vero cell-derived JCV/03 and rJ.03-G (Figure 7D, compare dashed and solid lines). As seen previously, growth of parental JCV/03 in C6/36 mosquito cells abolishes the replication lag seen with virus derived from mammalian cells (Figure 7D, compare solid and dashed red lines).

Similarly, passage of rJ.03-G in C6/36 cells allowed rapid growth in insect cells without an appreciable lag (Figure 7D, compare dashed and solid green lines). Thus, as anticipated, the rJ.03-G virus appears to behave identically to the parental JCV/03. In contrast, passage of rJ.03-S in C6/36 cells slightly improved replication in insect cells, but growth remained slower and less robust than the mosquito cell-derived viruses that encode 193G. Together, these results demonstrate that S193G governs the baseline replication phenotype of JCV/03, and that additional mutations arise during insect cell passage that independently enhance replication in C6/36 cells.

### 3.7. Multiple independent M segment mutations contribute to enhanced replication in C6/36 cells

In addition to the S193G substitution, sequencing identified numerous mutations that emerged during passage in C6/36 cells. All nonsynonymous mutations detected above the 5% cutoff are summarized in Supplemental Tables 6 and 7. Although the Series I and Series II experiments did not share any amino acid substitutions associated with adaptation, several M segment mutations displayed frequency changes that closely paralleled the replication phenotypes observed during the respective serial passage. In Series I, two nonsynonymous mutations in Gc, I1365V and L1394F, increased in frequency during passage in C6/36 cells and subsequently declined following passage in Vero cells, consistent with the reversion phenotype apparent in the final passage (Figure 8A). In Series II, two unique M segment mutations, P346A and H890Y, were associated with efficient replication in C6/36 cells and persisted after passage in Vero cells, consistent with the stable adaptation phenotype observed in the series (Figure 8B). Figure 8C summarizes the positions of these mutations within the JCV M segment polyprotein.

**Figure 8:**
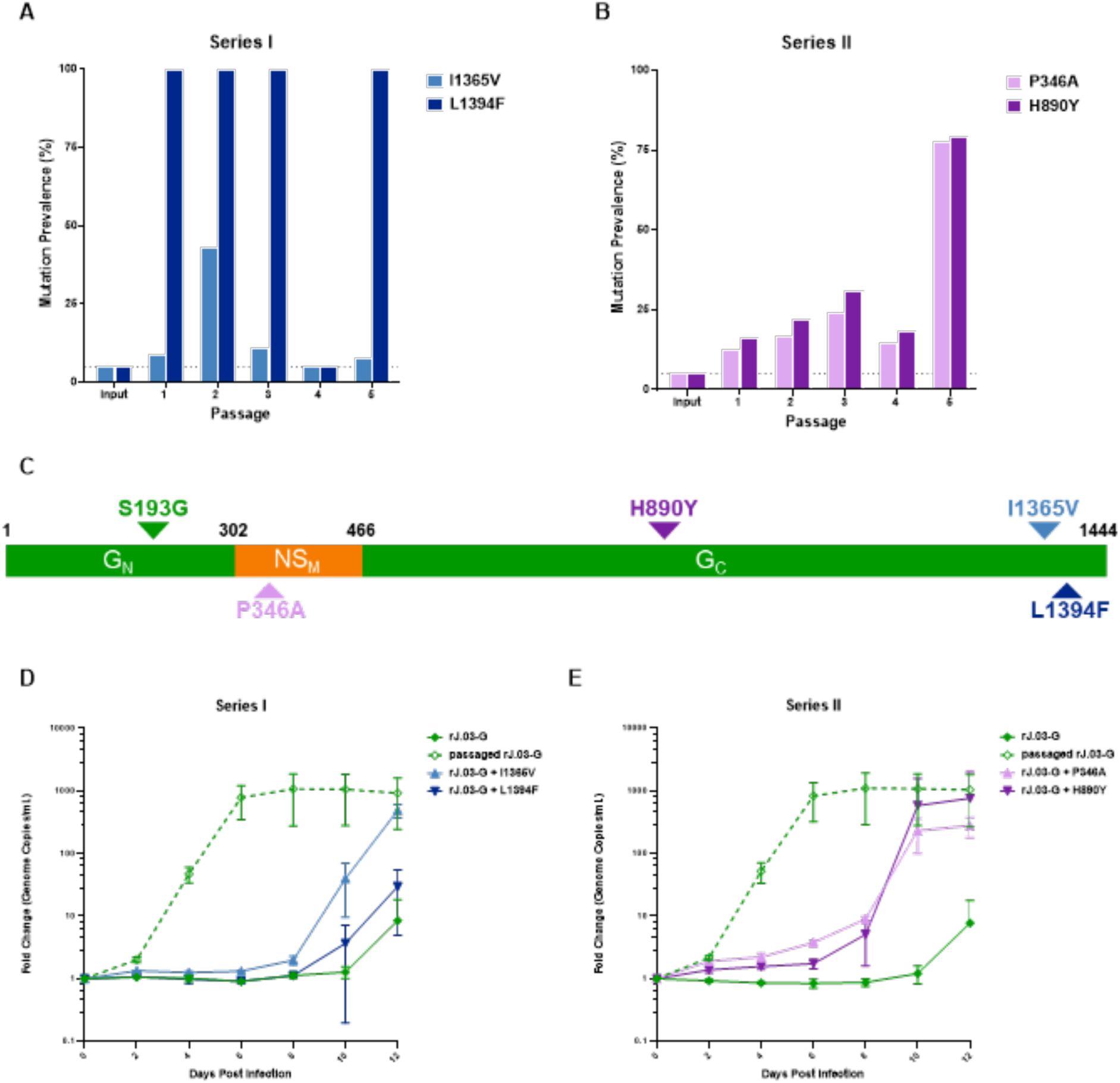
Multiple mutations across the M segment independently confer enhanced replication of JCV/03 in C6/36 cells. Mutation frequencies from Series I (A) and Series II (B) determined by Illumina sequencing of viral RNA isolated at the final time point of each passage. Only mutations that rose appreciably above background variation and whose frequencies correlated with the observed replication phenotypes. (C) Schematic representation of the JCV M segment polyprotein indicating Gn, NSm, and Gc regions. Positions of S193G and mutations identified during serial passage are indicated. Black numbers correspond to amino acid numbering based on the complete glycoprotein precursor. D,E) Multi-step growth curves in C6/36 cells comparing rJ.03-G to recombinant viruses carrying individual M segment mutations from Series I (D) and Series II (E). Supernatants were collected at the indicated time points and viral RNA was quantified by qPCR to determine genome copies per mL. Titers are expressed as fold change in genome copies per mL relative o input. Data represent the mean and SD of three biological replicates.

To determine whether these mutations contributed directly to enhanced replication in insect cells, each mutation was introduced individually into the M segment of rJ.03-G. After launch and amplification in Vero cells, these recombinants were evaluated for growth in mosquito cells and compared to Vero cell-derived rJ.03-G and C6/36-derived rJ.03-G. As observed previously, Vero-derived rJ.03-G displayed a prolonged lag phase, with viral titers remaining near baseline until day 10 post infection (Figures 8D and 8E, solid green lines). In contrast, C6/36-derived rJ.03-G transitioned to rapid replication by day 2 post infection and reached peak titers by day 6 (Figures 8D and 8E, dashed green lines). The Series I Gc mutations produced subtle changes in replication kinetics. rJ.03-G + I1365V entered rapid replication at day 8 post infection, approximately two days earlier than Vero-derived rJ.03-G and ultimately reached peak titers comparable to C6/36-derived rJ.03-G (Figure 8D, light blue line). rJ.03-G + L1394F also began increasing at day 8 post infection, but peak titers only modestly exceeded those of Vero-derived rJ.03-G (Figure 8D, dark blue line). The Series II mutations produced a stronger phenotype. Recombinant viruses carrying the P346A or H890Y mutations displayed detectable replication prior to day 8 post infection and then underwent rapid amplification beginning approximately day 8 (Figure 8E, purple lines). By day 10, both recombinants reached peak titers that were comparable to C6/36-derived rJ.03-G and remained stable through day 12. Thus, while all four mutations shortened the lag phase relative to Vero-derived rJ.03-G, they did not recreate the immediate, robust replication of C6/36-derived rJ.03-G.

In addition to P346A and H890Y, sequencing of Series II identified the NSm mutation W371R as a possible determinant of enhanced replication in C6/36 cells because its frequency increased during passages 1 and 2 in insect cells and subsequently declined during passage in Vero cells (Supplemental Table 7 and Supplemental Figure 3A). However, unlike P346A and H890Y, W371R was not detected in passage 5, suggesting it was not associated with the sustained adaptation phenotype observed in the growth curves. To directly test its contribution, recombinant rJ.03-G + W371R was generated and evaluated in C6/36 cells. The W371R encoding virus exhibited an abbreviated lag phase relative to both parental JCV/03 and Vero-derived rJ.03-G. Whereas parental JCV/03 and rJ.03-G transitioned to rapid replication at approximately days 6 and 4 post infection, respectively, rJ.03-G + W371R displayed exponential replication by day 2 post infection (Supplemental Figure 3B). Despite this accelerated onset of replication, rJ.03-G + W371R reached peak titers approximately 10-fold lower than parental JCV/03 and Vero-derived rJ.03-G. Peak titers were comparable to C6/36-derived rJ.03-G but, they were reached approximately two days later. Therefore, although W371R enhanced replication kinetics in C6/36 cells, this change alone did not reproduce the fully adapted phenotype.

We also performed sequence analysis of JCV/03 derived from serial passage at lower cell density (Supplemental Figure 2). This analysis identified a M352V substitution in NSm as another potential determinant of enhanced replication in C6/36 cells (Supplemental Figure 4A). Recombinant rJ.03-G + M352V displayed an intermediate phenotype between Vero-derived and C6/36-derived rJ.03-G. Following an approximately 8 day lag phase, rJ.03-G + M352V underwent rapid replication and reached peak titers by day 12 post infection (Supplemental Figure 4B). Peak titers were comparable to those of C6/36-derived rJ.03-G and greater than ten-fold higher than Vero-derived rJ.03-G. However, unlike C6/36-derived rJ.03-G, which entered rapid replication by day 2 and peaked by day 6, the M352V mutant retained a substantial replication delay. Thus, similar to the other M segment mutations evaluated in this study, M352V partially enhanced replication in C6/36 cells but was insufficient to fully reproduce the adapted phenotype.

Together, these data demonstrate that multiple independent mutations within the M segment can act to enhance JCV/03 replication in C6/36 cells. However, no single mutation fully recapitulated the rapid replication kinetics of C6/36-derived virus, suggesting that efficient adaptation likely reflects the combined effects of multiple mutations.

## 3. DISCUSSION

Jamestown Canyon virus (JCV) is unusual among arboviruses in its broad host and vector range, having been detected in more than 33 mosquito species, tabanids, and diverse mammalian hosts. (7, 8, 12, 36) To investigate viral determinants that may contribute to this ecological flexibility, we compared two genetically similar yet phenotypically distinct strains of JCV, JCV/61 and JCV/03. We identified a striking replication phenotype in *Aedes albopictus*-derived C6/36 cells in which JCV/03 exhibited a prolonged lag phase followed by abrupt, rapid replication. By contrast JCV/61 replicated immediately upon infection of mosquito cells. Importantly, a single passage of JCV/03 in C6/36 cells consistently eliminated the lag phase and conferred heritable enhancement of replication in mosquito cells. Sequencing of mosquito cell-adapted virus populations identified multiple mutations within the M segment that were selected during passage in C6/36 cells. However, we found that no individual mutation fully recapitulated the adapted phenotype, suggesting that multiple changes may be required to confer rapid and efficient replication in C6/36 cells. Overall, these results support a model whereby efficient replication in mosquito cells can arise through multiple evolutionary pathways.

A strength of this study was the development and application of a plasmid-based reverse genetics system for JCV/03. Concurrent efforts by Alyssa Evans and colleagues established a complementary reverse genetics platform for JCV/61 that has proven valuable for large-scale segment reassortment studies (37, 38). In contrast, the JCV/03 system described here was particularly well suited for targeted mutagenesis, allowing direct evaluation of specific amino acid substitutions. Together, these systems provide a foundation for future studies examining both segment-level and residue-level determinants of host adaptation in JCV.

When developing the reverse genetics system for JCV/03, we initially utilized the published reference genome from Bennett et al. which encoded a serine residue at position 193 (193S) of the Gn protein to construct the launch plasmids (27). Sequence analysis revealed that our parental stock and four independently derived biological clones differed from the reference sequence by encoding a glycine at this position (193G). Consequently, the recombinant virus generated from the published sequence encoded 193S (rJ.03-S), whereas our biological clone encoded 193G. In both Vero and C6/36 cells, rJ.03-S displayed replication kinetics distinct from parental JCV/03 and notably lacked the characteristic lag-burst phenotype in C6/36 cells that differentiated JCV/03 from the prototypic JCV/61. When glycine was introduced into the reverse genetic system, the resulting virus (rJ.03-G) recapitulated the parental phenotype in both mammalian and mosquito cells (Figures 7B and 7C). Although the mechanistic basis by which S193G contributes to the lag-burst phenotype remains unclear, residue 193 is clearly a critical determinant of this unusual replication behavior.

Analysis of orthobunyavirus sequences revealed that glycine position 193, as well as the surrounding region, is highly conserved in the genus. Moreover, no other JCV isolate encodes a serine at residue 193. The vast majority of orthobunyaviruses have a glycine at this position while the few that do not, typically encode an alanine. Given the overall sequence conservation in this region, it is unsurprising that the 193S recombinant virus showed impaired replication in both mammalian and insect cells. Inspection of the deposited reference JCV/03 sequence revealed that a single nucleotide misread could account for the serine to glycine exchange. These findings underscore the importance of verifying the exact sequence of recombinant viruses compared to parental viruses.

Using targeted mutagenesis, we tested seven substitutions identified in the serial passages distributed across Gn (S193G), NSm (P346A, M352V, W371R), and Gc (H890Y, I1365V, L1394F). All of the tested mutations enhanced JCV/03 replication in C6/36 cells to varying extents compared to parental virus. The location and identity of each mutation provide clues to their potential mechanism of action. Furthermore, analysis of publicly available sequence databases can help determine the likelihood that these mutations could emerge or persist in nature. Based on structural mapping of La Crosse virus glycoproteins, H890Y localizes to stalk subdomain 2 of Gc (39). Mutants lacking the stalk and head domains retained the ability to undergo low pH-triggered fusion, suggesting this region may serve a structural rather than pH-sensing role (40, 41). Substitution of histidine with tyrosine introduces a hydroxyl-containing aromatic side chain capable of altered hydrogen bonding, potentially influencing protein conformation or interactions. Neither the location nor biochemical properties of I1365V provide an obvious explanation for how this substitution enhances replication in C6/36 cells. However, because the mutation increased during passage in mosquito cells and declined following passage in Vero cells, the selective advantage conferred by I1365V appears to be host dependent. We found nearly 30 separate orthobunyavirus sequences with 1365V in the Bacterial and Viral Bioinformatics Resource Center (BVBRC), the most common residue at this position after I. Likewise, L1394F was not unique to our study, among the sequences on the BVBRC, leucine and phenylalanine were both common residues to find in this position. Further analysis is needed to determine whether viruses carrying 1365V and 1394F were recovered from mosquito or mammalian hosts.

Structurally, L1394F localizes immediately adjacent to a predicted transmembrane domain in a region analogous to the membrane-proximal external region (MPER) found in other viral fusion proteins (42–44). Studies have shown that residues within the MPER contribute to viral entry, intracellular trafficking, and membrane fusion, suggesting this region plays an important structural role during infection (45–47). The substitution of leucine with phenylalanine could therefore alter glycoprotein organization by strengthening intermolecular interactions through aromatic packing or by increasing membrane affinity near the viral envelope. Interestingly, although L1394F repeatedly reached fixation during serial passage, recombinant rJ.03-G + I1365V produced a more pronounced enhancement than rJ.03-G + L1394F (Figure 8D). This discrepancy suggests that mutation frequency alone may not directly reflect the magnitude of phenotypic contribution. Notably, I1365V and L1394F emerged together during serial passage, raising the possibility that these substitutions function cooperatively and that both may be required to fully reproduce the adapted phenotype. Determining the mechanistic basis for these phenotypes will require direct analysis of how individual and combined mutations glycoprotein fusogenicity and membrane interactions.

Unexpectedly, three of the mutations we identified and tested mapped to NSm. This finding was notable because NSm remains one of the least well-characterized proteins encoded by orthobunyaviruses. Moreover, with the exception of the N-terminal region, NSm is largely dispensable for replication in cultured mammalian and arthropod cells. P346A and M352V localized to NSm domain II, a non-hydrophobic predicted luminal domain. When engineered into rJ.03-G, both P346A and M352V enhanced JCV/03 replication in C6/36 cells, demonstrating that domain II tolerates sequence variation and that alterations within this region can improve replication without disrupting essential NSm functions. Both positions also appear highly conserved and are invariant within the California serogroup orthobunyaviruses including La Crosse, California encephalitis, Snowshoe Hare and Tahyna. Lastly, the W371R substitution localized to domain III of NSm, a canonical transmembrane domain implicated in membrane association and efficient viral replication (48, 49). In Bunyamwera, deletion of domain III substantially impairs replication (41). Replacement of a bulky hydrophobic tryptophan with a positively charged arginine in this domain might disrupt local membrane interactions and impair protein function. Nevertheless, recombinant rJ.03-G + W371R exhibited markedly accelerated replication in C6/36 cells (Supplemental Figure 3). The repeated emergence of NSm mutations during serial passage, together with their ability to enhance replication when tested individually, suggests that NSm contributes to mosquito cell adaptation through mechanisms that remain poorly understood.

In addition to the viral genetic alterations, we also observed that the insect cell culture conditions affected replication delay. The most notable was initial plating density which dramatically shortened the lag phase. It may be that higher cell densities permitted more efficient cell-cell spread thereby shortening the replication delay. Another possibility is that C6/36 cultures contain heterogeneous subpopulations of cells that differ in permissiveness to JCV/03 replication. Under this model, viral replication remains below the limit of detection until a sufficient number of permissive cells become available. Transcriptomic analysis of infected and uninfected C6/36 cells throughout the time course could reveal cellular pathways associated with the transition to rapid replication and generate new mechanistic hypotheses. More broadly, it remains unclear whether the lag-burst phenotype is unique to C6/36 cells or reflects a characteristic of mosquito-derived cultures. Future studies in additional insect cell lines are necessary to address this question and evaluate alternative explanations for the phenotype.

A longstanding theory in arbovirus evolution is that alternating replication between vertebrate and arthropod hosts constrains viral evolution (50–52). Under this trade-off hypothesis, mutations that improve fitness in one host are expected to reduce fitness in the alternate host, thereby limiting specialization and slowing long-term evolution. However, several studies have demonstrated that adaptation to one host does not necessarily impose a fitness cost in the other. For example, experimental evolution studies of dengue virus and alphaviruses, including Sindbis virus and Eastern equine encephalitis virus, have shown that some adaptive mutations can be neutral or even advantageous across hosts (50, 53). Similarly, the E1-A226V mutation in chikungunya virus enhanced replication and transmission in *Aedes albopictus* mosquitoes without compromising fitness in human hosts and is thought to have contributed to the virus’s global emergence (54). Our findings are broadly consistent with a more nuanced view of arbovirus evolution in which host alternation constrains some outcomes without completely preventing evolutionary innovation. In the first serial passage experiment adaptation to C6/36 cells produced little detectable effect on replication in Vero cells, yet the phenotype reverted following mammalian passage, suggesting that the adaptive changes were not stably maintained in the absence of mosquito cell-specific selection. In contrast, the second serial passage experiment demonstrated that adaptation acquired in mosquito cells could persist after passage through mammalian cells, although this persistence was accompanied by reduced replication in Vero cells. Together, these outcomes suggest that alternating-host replication acts not as an absolute evolutionary barrier but rather as a selective filter that may favor adaptive solutions compatible with both hosts.

Several limitations should be considered when interpreting these findings. First, our analysis focused exclusively on nonsynonymous mutations within coding regions and therefore did not assess the potential contributions of synonymous mutations or noncoding RNA elements. Second, all experiments were performed in relatively immunodeficient cell culture systems, as C6/36 cells lack a functional antiviral RNAi response and Vero cells cannot produce type I interferons (55, 56). Consequently, the mutations we assessed here likely arose from selective pressures other than evading host immunity. One selective pressure may be incubation temperature, as C6/36 and Vero cells are maintained at substantially different temperatures. Thus, it remains unclear whether the mutations selected during passage reflect adaptation to host-specific cellular factors, temperature, or both. Future studies in live mosquitoes and cell culture at non-native temperatures could help to distinguish among these possibilities. Finally, serial passage may promote accumulation of defective interfering (DI) particles that alter replication dynamics. Although normalization by genome copy number likely reduced the effective multiplicity of infection and partially mitigated these effects, the evolutionary pathways identified here should nevertheless be interpreted as models of laboratory adaptation rather than direct representations of viral evolution in nature.

Future studies aim to determine the biological relevance of the adaptive mutations identified here under more natural transmission conditions. Although all seven mutations enhanced replication in C6/36 cells, it remains unclear whether they confer similar advantages in live mosquito vectors. An important next step will also be to determine whether the mutations act independently or cooperatively. Because no individual mutation fully recapitulated the phenotype acquired during serial passage, combinations of mutations may be required to reproduce the rapid replication kinetics of mosquito cell-adapted virus.

In summary, this study identifies a previously unrecognized mosquito cell-specific replication phenotype in a Jamestown Canyon virus isolate and demonstrates that adaptation to mosquito cells can occur rapidly through multiple independent mutations within the M segment. The discovery that both glycoprotein and NSm mutations enhance replication highlights the diversity of genetic pathways available for mosquito-cell adaptation. More broadly, these findings establish a facile reverse genetics system for JCV/03 and provide a framework for dissecting the viral determinants that govern host-specific fitness and arbovirus evolution.

## Author Contributions

Conceptualization, PB and SD; Methodology, SD and TS; Investigation, SD, TS and PB; Resources, PB and SC; Writing—original draft preparation, SD; Writing—review and editing, PB, SC and SD; Supervision, PB and SC; Project administration, PB; Funding acquisition, PB and SC. All authors have read and agreed to the published version of the manuscript.

## Data Availability Statement

All data needed to evaluate the conclusions in the paper are included in the main paper and supplemental material. Reagents are available upon request from the corresponding authors.

## Acknowledgments

The authors thank Philip Armstrong and Theodore Andreadis of the Connecticut Agricultural Experiment Station for generously providing the JCV/03 stock used in this study. The authors also acknowledge the NIH Biodefense and Emerging Infections Research Resources Repository (BEI Resources) for supplying the JCV/61 strain and the NIH Bacterial and Viral Bioinformatics Resource Center (BV-BRC) for facilitating our genome work. Finally, the authors thank the Penn Medicine Translational Neuroscience Core for providing primary rat cortical neuron cultures and associated technical support.

