## Supplemental Figures for "Jamestown Canyon virus rapidly adapts to mosquito cells through multiple M segment mutations"

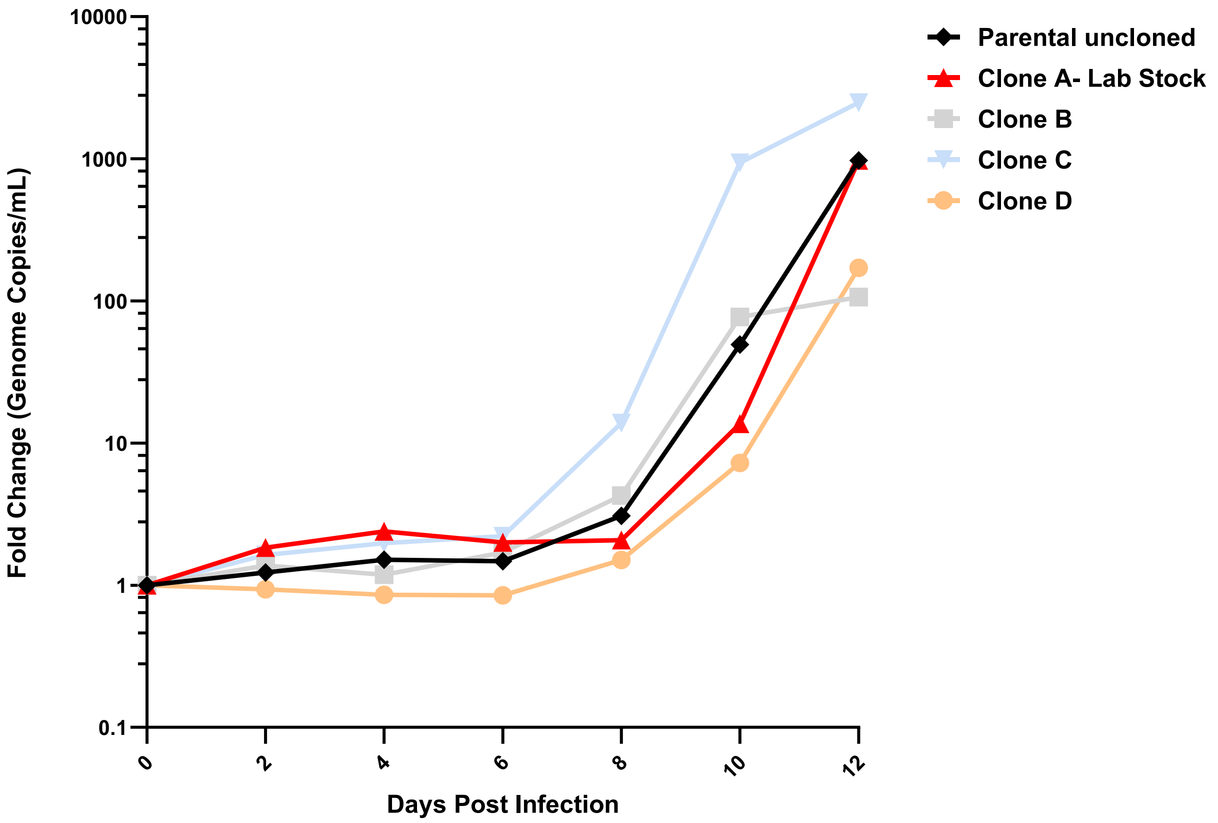


**Supplemental Figure 1. Parental, uncloned JCV/03 and biological clones of JCV/03 display the lag-burst growth phenotype in C6/36 cells.** C6/36 cells were infected at MOI 0.003 with parental, uncloned JCV/03 and four clones of JCV/03 generated after three rounds of terminal dilution. Supernatants were collected at the indicated time points and viral RNA was quantified by qPCR to determine genome copies per mL. Titers are expressed as fold change in genome copies per mL relative to input. One biological replicate per virus.

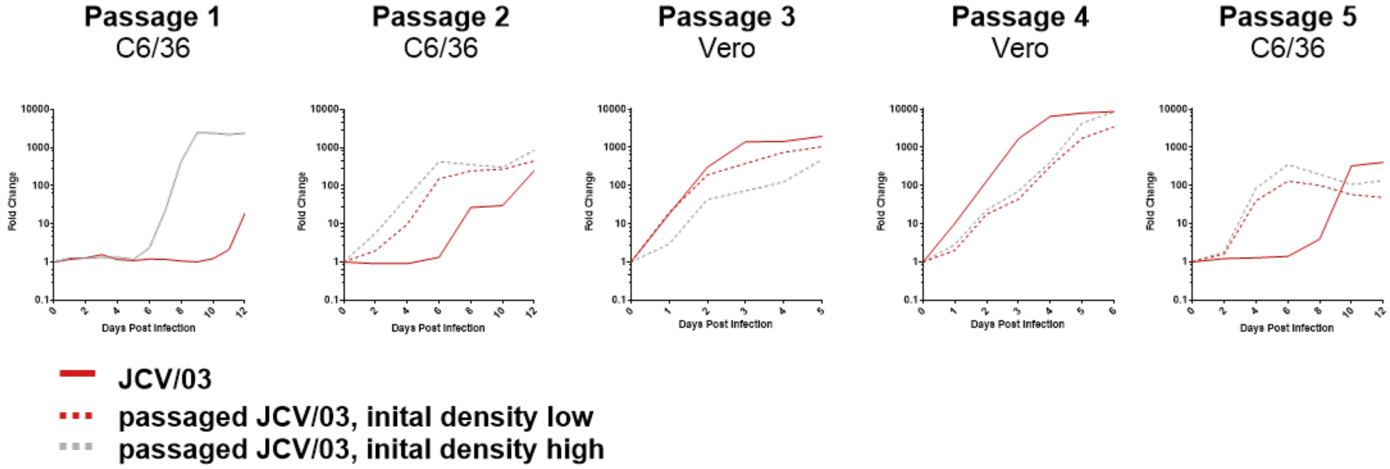


**Supplemental Figure 2: Initial plating density does not alter the long-term outcome of serial passage.** JCV/03 was serially passaged using low-density C6/36 cultures for passage 1, followed by serial passage as described for Figure 6. Cells used for passage 1 were plated at a density six-fold lower than the high-density condition. Virus was passaged twice in C6/36 cells (Passages 1 and 2), followed by two passages in Vero cells (Passages 3 and 4), and subsequently reinfected onto C6/36 cells (Passage 5). Cells were infected with 2 × $10^6$ genome copies of each virus and supernatants were collected daily. Viral RNA was quantified by qPCR and titers are expressed as fold change in genome copies per mL relative to input.

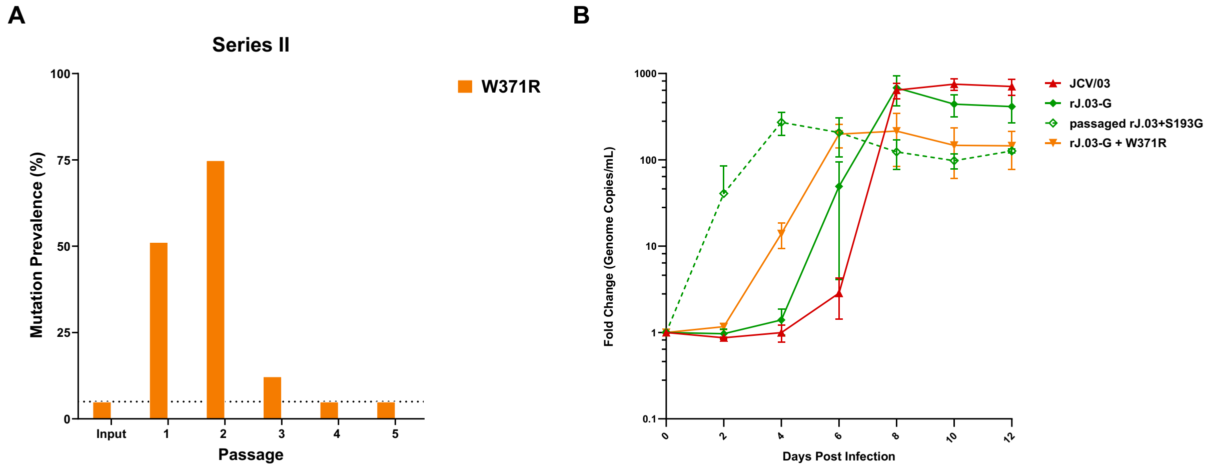


**Supplemental Figure 3: W371R enhances replication kinetics of rJ.03-G in C6/36 cells.** (A) The prevalence of the W371R mutation after serial passage as determined by Illumina sequencing of viral RNA isolated at the final time point of each passage. (B) C6/36 cells were infected with 2 × $10^6$ genome copies of each virus and supernatant was collected at the indicated timepoints. Viral RNA was quantified by qPCR and titers are expressed as fold-change in genome copies per mL. Data represent mean and SD of three biological replicates.

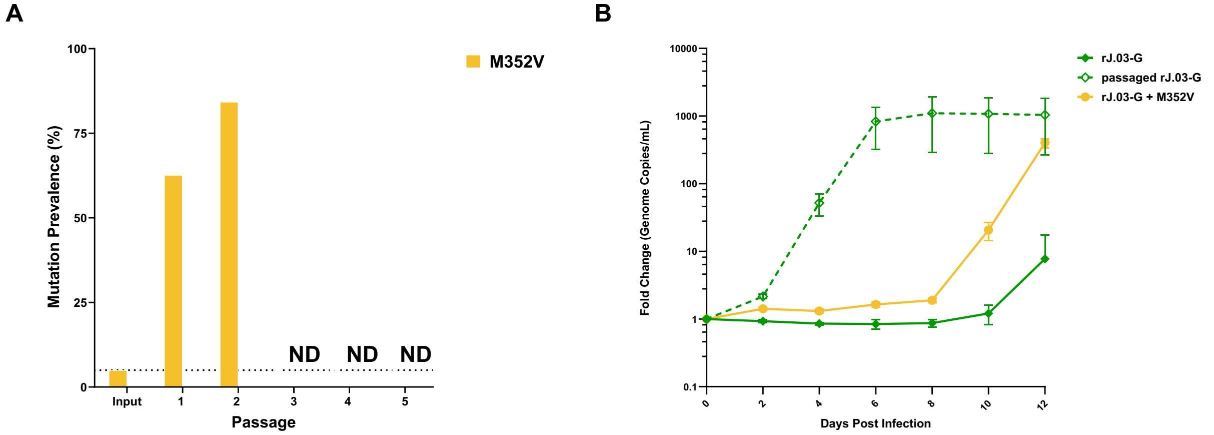


**Supplemental Figure 4: M352V enhances replication kinetics of rJ.03-G in C6/36 cells.** (A) The prevalence of the M352V mutation after serial passage as determined by Illumina sequencing of viral RNA isolated at the final time point of each passage. ND- not determined. (B) Multi-step growth curves in C6/36 cells comparing Vero-derived rJ.03-G, C6/36-derived rJ.03-G, and recombinant rJ.03-G + M352V. C6/36 cells were infected with $2 \times 10^6$ genome copies of each virus and supernatants were collected at the indicated time points. Viral RNA was quantified by qPCR and titers are expressed as fold change in genome copies per mL relative to input. Data represent mean and SD of three biological replicates.
