## Supplemental Tables for "Jamestown Canyon virus rapidly adapts to mosquito cells through multiple M segment mutations"

| Target | Forward Primer | Reverse Primer |
|---|---|---|
| JCV Small Segment (26) | TGACCAGTAGTGTACTCCAC | CAAGCAGTAGTGTGCTCCAC |
| JCV Medium Segment | AGTAGTGTGCTACCAAGTATATTTAAATGATTC | AGTAGTGTACTACCAAGTATAGAAAACGTTC |
| JCV Large Segment | AGTAGTGTGCTCCTATTTAC | AGTAGTGTACTCCTATTTACAAAACTTAC |

**Supplemental Table 1:** Primers used for sequencing biological clones and verifying point mutations

| Virus & Segment | Nucleotide | | | Amino Acid | | |
|---|---|---|---|---|---|---|
| | Position | Reference | Sequencing | Position | Reference | Sequencing |
| JCV/61 S | 9 | A | G* | N/A | | |
| JCV/61 M | 1-25 | | N/D | | | |
| JCV/61 M | 328 | T | C | 92 | N | N |
| JCV/61 M | 759 | C | T | 236 | T | I |
| JCV/61 M | 885 | A | C | 278 | N | T |
| JCV/61 M | 1067 | C | G | 339 | L | V |
| JCV/61 M | 1140 | C | V | 363 | T | I |
| JCV/61 M | 1174 | T | A | 374 | F | L |
| JCV/61 M | 1176-1177 | GC | TG | 375 | C | L |
| JCV/61 M | 1240 | G | A | 396 | A | A |
| JCV/61 M | 1534 | C | T | 494 | T | T |
| JCV/61 M | 1619 | G | A | 523 | G | R |
| JCV/61 M | 1738 | C | T | 562 | A | A |
| JCV/61 M | 1783 | A | C | 577 | K | N |
| JCV/61 M | 1846 | T | C | 598 | V | V |
| JCV/61 M | 2051 | G | A | 667 | E | K |
| JCV/61 M | 2466 | G | A | 805 | G | D |
| JCV/61 M | 3130 | G | A | 1026 | K | K |
| JCV/61 M | 3190 | A | G | 1046 | G | G |
| JCV/61 M | 3852 | G | A | 1267 | S | N |
| JCV/61 M | 3931 | G | A | 1293 | E | E |
| JCV/61 M | 4070 | C | A | 1340 | H | N |
| JCV/61 M | 4485-4510 | | N/D | | | |
| JCV/61 L | 1-26 | | N/D | | | |
| JCV/61 L | 5013 | A | G | 1652 | L | L |
| JCV/61 L | 6631 | G | A | 2192 | D | N |
| JCV/61 L | 6900-6960 | | N/D | | | |
| JCV/03 S | 9 | A | G* | | | |
| JCV/03 M | 1-24 | | N/D | | | |
| JCV/03 M | 629 | A | G | 193 | S | G |
| JCV/03 M | 4484-4510 | | N/D | | | |
| JCV/03 L | 1-22 | | N/D | | | |
| JCV/03 L | 6935-6957 | | N/D | | | |

**Supplemental Table 2: Sequence comparison between prototype JCV clones and the designated reference sequence.** Sequences derived from the prototype strain of JCV/61 and JCV/03 were aligned to the corresponding reference genome. JCV/61 (Small: U12796.1, Medium: U88058.1, Large: NC_043559.1) JCV/03 (Small: HM007353.1, Medium: HM007354.1, Large: HM007355.1) * represents a mutation introduced by the primer used for sequencing. N/D – not determined at this time. N/A – not available because the mutation occurred in a non-coding region and therefore, there was no associated amino acid.

| Target | Forward Primer | Reverse Primer |
|---|---|---|
| JCV/03 N | AAACACGATGATAATATGGGAGATTTGGT TTTCTATGATG | GATCTCAGTGGTCTATCAAGGCAGTTTTA CACCG |
| pIRES-polyA vector | TAGACCACTGAGATCTGACTGAAA | ATTATCATCGTGTTTTTCAAAGGAA |

**Supplemental Table 3: Primers used for constructing JCV/03-pN expression plasmid**

| Target | Forward Primer | Reverse Primer |
|---|---|---|
| JCV N | CCTGCCACTCTCCAAACTTAG | TTCCTTAATGCAGCAAAAGCC |
| C6/36 Actin (29) | CACGAACTGGGACGATATGGA | CGGTTAGCCTTCGGGTTCAG |

**Supplemental Table 4: qPCR primers**

| Mutation | Forward Primer | Reverse Primer |
|---|---|---|
| S193G | TACTCCCTGGTTCAATAGCTAACTCTATAT GCCA | TTGAACCAGGGAGTATTGTTCTGTGTAAA AAC |
| P246A | TGATTTAGCCGATGATATGGTACAAATGGC | GTACCATATCATCGGCTAAATCATCTAGA GTG |
| M352V | GATATGGTACAAGTGGCTGAGAGAGTGAA CATTTAC | CACTCTCTCAGCCACTTGTACCATATCAT C |
| W371R | GTGTTACACGGGCTTTCTTGCTACTTG | AAGCCCGTGTAACACCATAATTTATAATG AT |
| H890Y | ATGTGGGATATTTTTGCCTTAGTCCCAGAT GC | AAAAATATCCCACATCTTGATCTTTATCAT AA |
| I1365V | CTGCATATGTCAAAGAGAAAGATGAGAGAT GT | CTTTGACATATGCAGTTTGGTCTACTGGT GCA |
| L1394F | CCTTTTAAAAATTTTTTCGGCTCTTATATTG GTATATTTTATACAGGC | GCCGAAAAAATTTTTAAAAGGCTCTAATA TAACTTGAAATCC |

**Supplemental Table 5. Primers used to introduce point mutations in the JCV/03 pM launch plasmid**

| Gene | Mutation | Input | Pass 1-C6/36 | Pass 2-C6/36 | Pass 3-Vero | Pass 4-Vero | Pass 5-C6/36 |
|---|---|---|---|---|---|---|---|
| Polymerase | A1504T | | 17.7% | 15.0% | 12.3% | 5.3% | |
| Polymerase | F1962L | | | | 7.0% | 7.0% | 14.0% |
| NSm | D347N | | 13.7% | | 7.4% | 5.6% | |
| NSm | L378P | | | | 6.4% | 50.8% | |
| NSm | Y395H | | 44.3% | 44.3% | 70.5% | 38.0% | 66.0% |
| NSm | M404V | | 6.3% | | | | |
| NsM/Gc | K467E | | | 17.0% | | | |
| Gc | D514N | | | | | 38.6% | |
| Gc | R590W | | | | | | 8.9% |
| Gc | E625G | 24.7% | 25.4% | 5.7% | 16.1% | 53.4% | 11.1% |
| Gc | H890L | | 9.3% | 14.6% | | | |
| Gc | T1233I | | | | | | 73.1% |
| Gc | P1352S | 23.1% | | | | | |
| Gc | I1365V | | 8.8% | 43.1% | 11.1% | | 7.6% |
| Gc | 1393_1394insF | | 9.4% | 15.6% | 15.8% | 7.7% | 17.0% |
| Gc | L1394F | | 100.0% | 100.0% | 100.0% | | 100.0% |
| Gc | F1402Y | | 47.2% | 41.8% | 68.4% | 42.0% | 68.0% |

**Supplemental Table 6: Combined sequencing data for all non-synonymous mutations discovered in Series I.** Mutation frequencies from Series I determined by Illumina sequencing of viral RNA isolated at the final time point of each passage. Mutations highlighted in green were chosen for follow up in Figure 8. Amino acid numbering based on the open reading frame in the reference genome for the polymerase (large segment), polyprotein (medium segment), and nucleocapsid (small segment). Accession numbers for JCV/03, Small: HM007353.1, Medium: HM007354.1, Large: HM007355.1 Mutation S193G not shown. Minimum variant frequency cutoff of 5%.

| Gene | Mutation | Input | Pass 1-C6/36 | Pass 2-C6/36 | Pass 3-Vero | Pass 4-Vero | Pass 5-C6/36 |
|---|---|---|---|---|---|---|---|
| Polymerase | V1656I | | | | | | 6.6% |
| Polymerase | I1706T | | | 8.2% | | | |
| Gn | A174V | | | 7.9% | | | |
| NSm | L342P | | 9.8% | | | 7.3% | |
| NSm | P346A | | 12.6% | 16.8% | 24.0% | 14.7% | 77.4% |
| NSm | W371R | | 51.2% | 74.9% | 12.3% | | |
| NSm | G377D | | 7.0% | | | | |
| NSm | C402Y | | 11.7% | | 14.8% | 6.7% | |
| Gc | I540L | | | | 32.7% | 62.1% | 17.2% |
| Gc | H554R | | | | | 6.7% | |
| Gc | E625G | 24.7% | 10.5% | | | 6.4% | |
| Gc | T628I | | | | | | 34.7% |
| Gc | H890Y | | 16.0% | 22.0% | 30.7% | 18.0% | 79.0% |
| Gc | H890L | | | | | | |
| Gc | T1233I | | | | | | 15.6% |
| Gc | P1352S | 23.1% | | | | | |
| Gc | Y1364H | | | 71.2% | | | |
| Gc | S1397F | | | | | | 63.9% |
| Gc | G1400S | | | | 5.8% | 9.1% | 9.4% |
| N/NSs | G22G/ G16V | | | | | 7.9% | |

**Supplemental Table 7: Combined sequencing data for all non-synonymous mutations discovered in Series II.** Mutation frequencies from Series II determined by Illumina sequencing of viral RNA isolated at the final time point of each passage. Mutations highlighted in green were chosen for follow up in Figure 8. Amino acid numbering based on the open reading frame in the reference genome for the polymerase (large segment), polyprotein (medium segment), and nucleocapsid (small segment). Accession numbers for JCV/03, Small: HM007353.1, Medium: HM007354.1, Large: HM007355.1 Mutation S193G not shown. Minimum variant frequency cutoff of 5%.
